# Screening for Pea Seed-borne Mosaic Virus Resistance (PSbMV) in Lentil and Development of Associated KASP Markers

**DOI:** 10.64898/2026.10.09.758073

**Authors:** Giri Raj Paudel, Dinesh Babu Paudel, Ana Priscilla Montenegro Alonso, Kirstin E. Bett, Sean M. Prager

## Abstract

Lentil (*Lens culinaris* M.) is one of the most important legume crops grown worldwide, valued for its high nutritional content and its role in improving soil health. Pea seed-borne mosaic virus (PSbMV) poses a significant threat to lentil production worldwide, highlighting the need to identify resistant germplasm and incorporate the trait into breeding programs for sustainable disease management. Although lentil lines carrying recessive resistance to PSbMV have been reported, the underlying resistance genes have not been characterized in detail, and no resistance-associated molecular markers are currently available. In this study, a diversity panel of 86 lentil lines was screened using rub inoculation under controlled conditions, followed by phenotyping through quantitative PCR assays. Virus titer in the inoculated and upper systemically infected leaves was measured and normalized with the expression of host reference gene as an internal control. Inoculated leaves showed higher virus titers, but titers in systemically infected leaves were low in some, but not all, lines, including the previously reported lines carrying the recessive gene *sbv*, which grouped as resistant lines. In pea, certain amino acid substitutions in eIF4E are associated with recessive resistance to PSbMV. To test if the lentil homolog acts in similar fashion, cDNAs from selected resistant and susceptible lines were sequenced and analyzed. Consistent with pea, several amino acid substitutions in lentil eIF4E were associated with resistance. Nucleotide polymorphisms underlying these amino acid substitutions were used to develop Kompetitive Allele-Specific PCR (KASP) primer sets, which effectively distinguished resistant from susceptible lentil lines.

## Introduction

Pea seed-borne mosaic virus (PSbMV, *Potyvirus pisumsemenportati*, Family *Potyviridae*) naturally infects lentil and other major cool-season legumes (Khetarpal and Maury 1987). PSbMV is identified as one of the major threats in major lentil-producing regions (Coutts et al. 2008; Mabrouk and Mansour 1998; Van Leur et al. 2025), including regions in the Pacific Northwest and Western Canada (Beck et al. 2018; Hampton and Muehlbauer 1977; Paudel et al. 2025). The symptoms of PSbMV in lentils include narrow and curled leaves, leaf mottling or yellowing, dying shoot tips, and misshapen, aborted, or undersized pods (Coutts et al. 2008; Hampton and Muehlbauer 1977). The development of symptoms and yield reduction are shown to depend on environmental factors, the stage of lentils at the time of virus inoculation, and host and viral genotypes (Coutts et al. 2008; Van Leur et al. 2026). In some genotypes, yield losses above 60% have been recorded (Mabrouk and Mansour 1998). PSbMV is vertically transmitted via seed with a transmission rate of up to 44% in some lentil genotypes (Hampton and Muehlbauer 1977). Field transmission of the virus mainly occurs in a nonpersistent manner by more than 20 aphid species (Khetarpal and Maury 1987). This dual transmission route makes PSbMV a serious threat to lentil production in different regions of the world (Rashed et al. 2018; Van Leur et al. 2026) and underscores the need for identifying naturally occurring resistance to the virus.

A search for natural resistance against PSbMV identified translation initiation factor eIF4E or its isoforms eIF(iso)4E-associated resistances in pea (Gao et al. 2004a; Gilpin et al. 1997; Provvidenti 1987). One of the best-characterized recessive resistances against PSbMV is governed by the *sbm* allele associated with the *eIF4E* on pea chromosome VI, which also confers resistance against several additional potyviruses (Andrade et al. 2009; Gao et al. 2004a; Gao et al. 2004b; Gilpin et al. 1997; Provvidenti 1987; Provvidenti and Hampton 1993). Comparative analysis of the eIF4E protein between susceptible and resistant pea lines revealed differences in a few amino acid residues (Ashby et al. 2011; Gao et al. 2004a; Konečná et al. 2014; Smýkal et al. 2010).

Kompetitive allele-specific PCR (KASP) genotyping provides a rapid, reproducible, and cost-effective alternative for screening large germplasm collections. It enables efficient discrimination of specific DNA sequence variants, including single nucleotide polymorphisms and small insertions/deletions. Nucleotide polymorphisms in eIF4E associated with resistance have been exploited to develop KASP markers for the rapid detection of PSbMV resistance in pea lines (Swisher Grimm and Porter 2020) and clover yellow vein virus (*Potyvirus trifolii*) resistance in common bean lines (Hart and Griffiths 2013). This supports the feasibility of a similar strategy in lentil.

Similar to recessive resistance in pea, four lentil lines (PI 212610, PI 251786, PI 297745, and PI 368648) with the recessive resistance gene *sbv,* have been reported in the literature. These lines showed no visible symptoms after virus inoculation (Haddad et al. 1978). However, these early observations mostly focused on the development of visual symptoms and did not further characterize the genetic basis or the molecular mechanisms underlying PSbMV resistance in lentil.

In this study, a lentil diversity panel was screened for PSbMV resistance using quantitative PCR (qPCR), and several resistant lentil lines with lower virus titers in the systemically infected leaves were identified. These lentil lines with low virus titer include previously reported lines with the *sbv* resistance gene. To assess the presence of eIF4E-associated recessive resistance in lentil, cDNAs from resistant and susceptible lentil lines were sequenced and compared with pea homologs to identify amino acid substitutions potentially associated with resistance or susceptibility. Single nucleotide polymorphisms corresponding to such substitutions were then targeted for KASP marker development, with the goal of distinguishing resistant and susceptible genotypes across the lentil diversity panel. This study provides the first evidence of eIF4E-associated PSbMV resistance in lentil and provides associated KASP markers that helps in marker-assisted selection and accelerate breeding for potyvirus resistance.

## Materials and Methods

### Plant growth conditions and virus inoculation

PSbMV resistance screening was carried out on a subset (83 out of 324) of a Lentil Diversity Panel (LDP), which includes lines sourced from various regions of the world (Haile et al. 2020). An additional three lines (PI 251786, PI 297745, and PI 368648), carrying the recessive resistance gene (*sbv*) against the PSbMV (Haddad et al. 1978), which are not the part of LDP were included in the study. Plants were grown individually in 12.7×12.7×11.4 cm pots filled with PRO-MIX^®^ BX (Premier Tech) growing medium in controlled growth chambers set to 22°C with a photoperiod of 16 hours of light and 8 hours of dark. PSbMV inocula were maintained in pea (CDC Meadow) plants and used for virus inoculation experiments as described by (Paudel et al. 2025). In short, systemically infected young leaves of pea were ground in a sterilized mortar and pestle, mixed with 5 times the volume of buffer (0.01 M phosphate buffer) and a small amount (∼1gm) of Carborundum, and rub-inoculated onto the lower fully expanded leaves of 14-day-old lentil plants. Due to logistical constraints, lines were evaluated in five batches, all maintained under identical environmental and agronomic conditions. Within each batch, both virus-inoculated plants and non-inoculated plants (from the same cohort as controls) were replicated three times.

### RNA extraction, reverse transcription, and genomic DNA extraction

Leaf tissue was collected 14 days post-inoculation (dpi) from inoculated and upper systemic leaves separately (Fig. 1) and placed into 1.5 mL tubes. Samples were either processed immediately for RNA extraction or stored at -80°C until RNA extraction. Total RNA was extracted from 100 mg of leaf tissue using TRIzol (ThermoFisher) reagent following the protocol described by the manufacturer, with minor modifications similar to those described in (Paudel et al. 2025). The extracted RNA pellet was then dissolved in 50 μL of RNase-free water, incubated at 60°C for 10 min in a water bath, snap-cooled on ice, and stored at -80°C for long-term preservation. Reverse transcription was performed using the First Strand cDNA Synthesis Kit (New England Biolabs, NEB) with oligo(dT)23 VN primers following the manufacturer’s quick protocol (www.neb.com/E6560). Genomic DNA was extracted from systemic leaves of one representative plant per line for 25 randomly selected lines, using a separate ∼100 mg leaf tissue sample collected at the same time as the sample taken for RNA extraction. DNA was extracted using the DNeasy Plant Mini Kit (Qiagen, USA) according to the manufacturer’s protocol (http://www.qiagen.com/HB-1166).

**Fig. 1.**
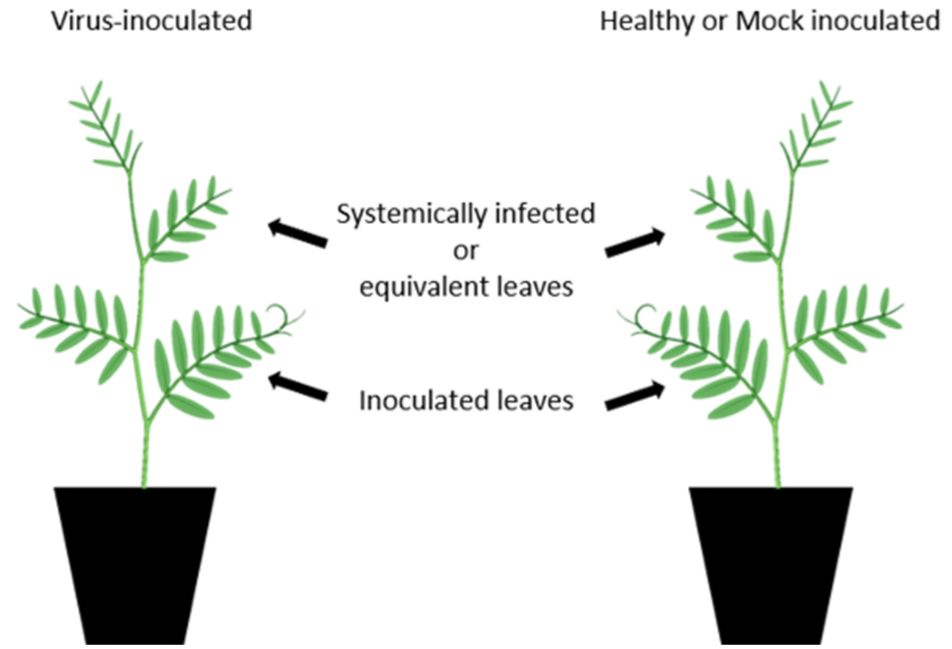
Diagrammatic representation of the leaflet positions used for inoculation and tissue sampling in lentil plants.

### PCR and qPCR conditions

PCR assays were performed from the cDNA using 2X FroggaBio Master Mix (FroggaBio Inc.) according to the manufacturer’s recommended protocol for a 20 μL reaction. The primer information used in this study to amplify PSbMV and lentil eukaryotic translation elongation factor 1 alpha (*LceEF1α*, as lentil reference) is listed in Table 1. Amplification was performed on an Applied Biosystems™ ProFlex™ PCR System at an annealing temperature of 57°C. The extension time was adjusted based on the expected amplicon size, at approximately 1 minute per kilobase of product as suggested for the polymerase mix. The final PCR products were run on an agarose gel and visualized using the iBright™1500 Imaging System (Thermo Fisher Scientific).

**Table 1:**
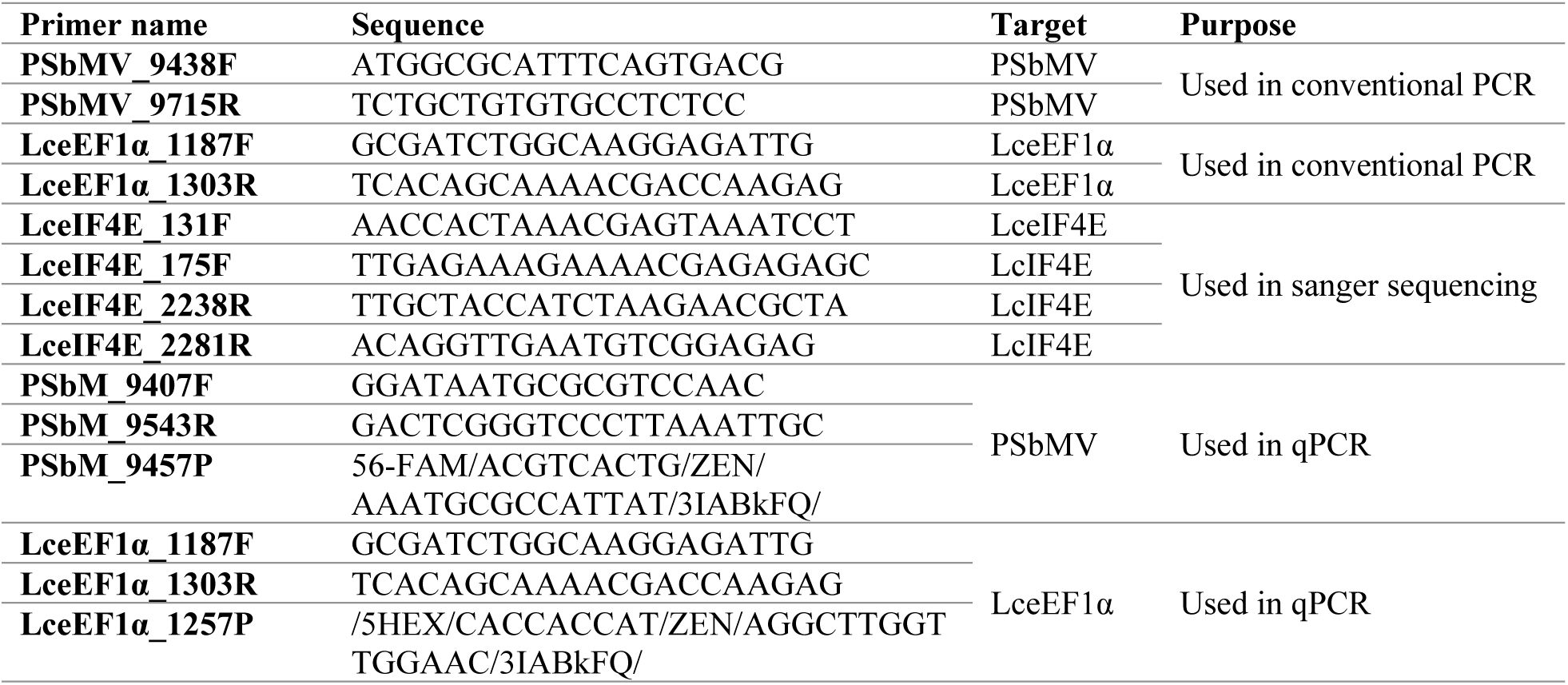
Primers sequences (5’-3’) and their corresponding reference targets, PSbMV (Pea seed-borne mosaic virus), lentil reference gene (LceEf1α), and LceIF4E (lentil translation initiation factor 4E). Forward primers are denoted with (F), reverse primers with (R), and probes with (P) at the end of their names.

PSbMV-specific primers were used to assess the presence of the virus in the inoculated leaves to evaluate the susceptibility of lentil lines to local viral replication, and in upper, non-inoculated leaves to determine the presence of systemic movement and subsequent viral multiplication. Comparing the virus and reference gene bands on the gel from conventional PCR, the virus titer cannot be quantified, and comparison among lentil lines is difficult. Since conventional PCR is not suitable for accurate quantification, a multiplex qPCR assay was performed to simultaneously measure PSbMV transcripts and a plant reference gene (*LceEF1α*) transcript in each sample, where a clear comparison of virus titer can be made among lentil lines by quantifying both virus and reference genes within the same sample.

Multiplex qPCR targeting the PSbMV and *LceEF1α* was performed as previously described (Paudel et al. 2025). The FAM-labelled PrimeTime qPCR assay (Integrated DNA Technologies, Inc., IDT) for PSbMV was used as described in that study (Table 1). A new HEX-labelled PrimeTime qPCR assay (IDT) targeting *LceEF1α* (NCBI accession number MF977709.1) was designed (Table 1) similarly to that for pea (Paudel et al. 2025). For the qPCR, lentil cDNA, TaqMan™ Fast Advanced Master Mix (Thermo Fisher Scientific), FAM (PSbMV) and HEX (*LceEF1α*) -labeled PrimeTime qPCR assays were mixed in a total reaction volume of 20 μL and run on a QuantStudio™ 3 Real-Time PCR System (Thermo Fisher Scientific, USA). Cycle thresholds for the lentil and PSbMV for each sample were exported and further analysis was performed in MS Excel. The ΔCT values were calculated by subtracting the Cycle threshold (CT) for *LceEF1α* (CTL) from the PSbMV CT (CTP), and the average ΔCT values from three replicates for each line were presented (Supplementary file 1).

### Sanger sequencing of lentil *eIF4E*

The pea *eIF4E* nucleotide sequence from the resistant pea line PI 269818 (Gao et al. 2004b) was retrieved from the NCBI database (Accession no: AY611425) and used to search for lentil homologs in the Phytozome 14 database (https://phytozome-next.jgi.doe.gov/). A gene (Lcu.2RBY.2g006900) was identified in the CDC Redberry lentil reference genome V2.0 (Ramsay et al. 2021) on Lcu.2RBY.Chr2:15192241-15194998 (reverse strand), as a homolog of the pea *eIF4E*. The complete genomic sequence of the lentil homolog (*LceIF4E*) was downloaded, and forward and reverse primers (Table 1) targeting the coding region were designed from the 5’ and 3’ ends of the genomic sequence using SnapGene (v8.2.2) (from Dotmatics; available at snapgene.com).

Lentil cDNAs from selected susceptible and resistant lines were PCR-amplified with a combination of *LceIF4E*-specific forward and reverse primers (Table 1) with Q5® High-Fidelity DNA polymerase (NEB), following the manufacturer’s protocol. The PCR products were then either gel-purified using the QIAquick Gel Extraction Kit (Qiagen) or cleaned using the Exo-CIP™ Rapid PCR Cleanup Kit (NEB), following the manufacturers’ standard protocols. These cleaned PCR products were then Sanger sequenced using one or more *eIF4E*-specific forward and reverse primers (Table 1). Sanger sequencing reads were processed to remove low-quality ends manually using SnapGene (v8.2.2). High-quality reads from each line were assembled, and the contigs were then compared with the CDC Redberry sequence to deduce the translation frame and amino acid sequence. Partial nucleotide sequences of lentil *eIF4E* for CDC Greenstar (PZ761244), CDC Impower (PZ761245), CDC Maxim (PZ761246), Crimson AGL (PZ761247), PI 297745 (PZ761248), PI 368648 (PZ761249), PI 518734 (PZ761250), CN105862 (PZ761251), ILL 213_AGL (PZ761252), ILL 2194 (PZ761253), PI 329167 (PZ761254), PI 472213 (PZ761255) were generated and submitted to the NCBI database (Accession numbers #PZ761244-PZ761255).

### Protein homology modeling

The amino acid sequences of eIF4E from two lentil lines (PI 297745 and ILL 213_AGL) were individually submitted to the SWISS-MODEL online server (Waterhouse et al. 2018) to generate 3D protein structure models. Homology modeling was performed using the pea eIF4E crystal structure (PDB accession code: 2WMC, chain H) (Ashby et al. 2011) as the template. One structural model was generated for each lentil eIF4E protein. The resulting 3D models were visualized and analyzed using ChimeraX (Meng et al. 2023).

### KASP marker design, assay, and validation

Available nucleotide sequences of the lentil eIF4E were aligned, and polymorphisms between the resistant and susceptible lines were identified using SnapGene (v8.2.2). Three KASP by Design Primer Mixes (LceIF4E-K1, LceIF4E-K2, and LceIF4E-K3; Table 2) were designed to target the three major single nucleotide polymorphisms that consistently differed between susceptible and resistant lentil lines. The KASP marker assays were performed using the protocol outlined by LGC Limited (http://www.lgcgenomics.com). Briefly, the 10.14 µL KASP assay mix was prepared with 0.14 µL KASP by Design Primer Mix, 5 µL KASP-TF Master Mix, and 5 µL DNA per sample. Amplification and visualization were performed on a QuantStudio 3™ Real-Time PCR System (Thermo Fisher Scientific, USA) with the following thermocycling conditions: a pre-read stage of 1 min at 30 °C, followed by a hold stage at 94 °C for 1 min (ramp rate 1.6 °C/s), followed by a touchdown phase consisting of 10 cycles of 94°C for 20 s, then 61°C for 1 min, decreasing by 0.6°C per cycle until reaching the target annealing temperature of 55°C. The touchdown stage was followed by 26 amplification cycles, each consisting of 94°C for 20 s and 55°C for 1 min. A post-read stage was performed at 30°C for 1 min to stabilize the endpoint fluorescence. To improve cluster separation, a recycling run was performed after the initial assay. The recycling program consisted of three cycles of 94°C for 20 s and 57°C for 1 min, followed by a final plate read at 30°C for 1 min. Genotype calls were generated with the QuantStudio™ Design and Analysis Software (Thermo Fisher Scientific) by assigning samples to allele clusters based on endpoint fluorescence values.

**Table 2:**
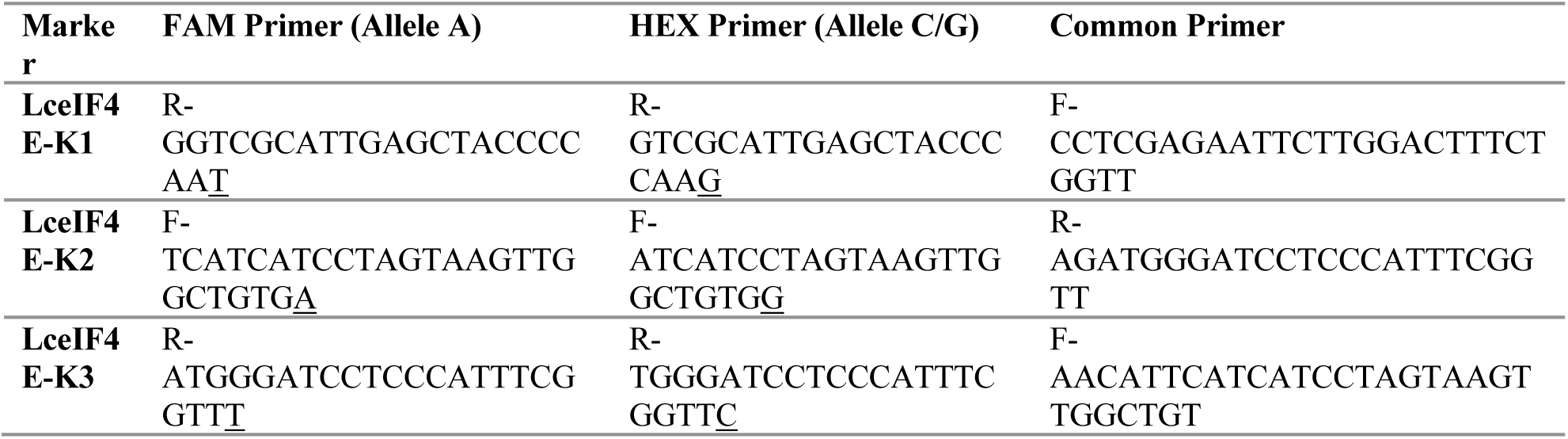
Primers used in the KASP assay. Forward primers in response to the eIF4E coding sequences are denoted with (F), and reverse primers with (R). The single-nucleotide polymorphisms targeted by the KASP markers are underlined. LceIF4E-K1, LceIF4E-K2, and LceIF4E-K3 markers target single-nucleotide polymorphisms observed at C221A, G319A, and G352A nucleotide positions of CDC Redberry eIF4E, respectively.

### Screening lentil lines for virus resistance using a PCR-based approach

A total of 86 diverse lentil lines, including four (PI 212610, PI 251786, PI 297745, and PI 368648) previously reported PSbMV-resistant lines (Haddad et al. 1978) were inoculated with PSbMV and evaluated for disease development. Using PSbMV-specific primers, expected amplicons in both inoculated and systemically infected leaves were consistently detected, including in the lines previously reported to be immune based on visual symptoms (Fig. 2). This strongly suggests that the replication and systemic movement of PSbMV can occur in all lentil lines, even in asymptomatic plants classified as resistant. Furthermore, characteristic symptoms in the lentil lines were observed at 30 days post-inoculation. While there were some lines that did not show any symptoms (PI 299165 AGL, PI 368648) at 30 days post-inoculation, others exhibited leaf chlorosis, reduced leaflet size, dwarfing, and leaflet distortion (Fig. 3).

**Fig. 2.**
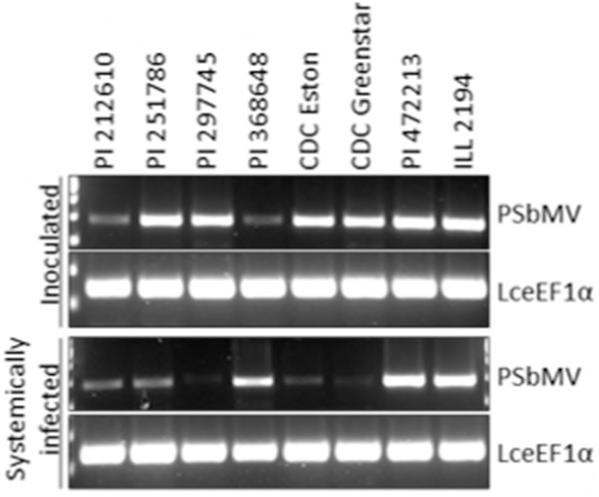
Gel image showing the PCR products from inoculated (top) and systemic leaves (bottom) across 8 representative lentil lines at 30 days post-inoculation with PSbMV. The PSbMV and lentil cDNA were amplified with PSbM_9438F/PSbM_9715R and LceEF1α _1187F/LceEF1α _1303R primer pairs, respectively, using the cDNA obtained from inoculated (upper panel) and systemically infected (lower panel) leaves.

**Fig. 3.**
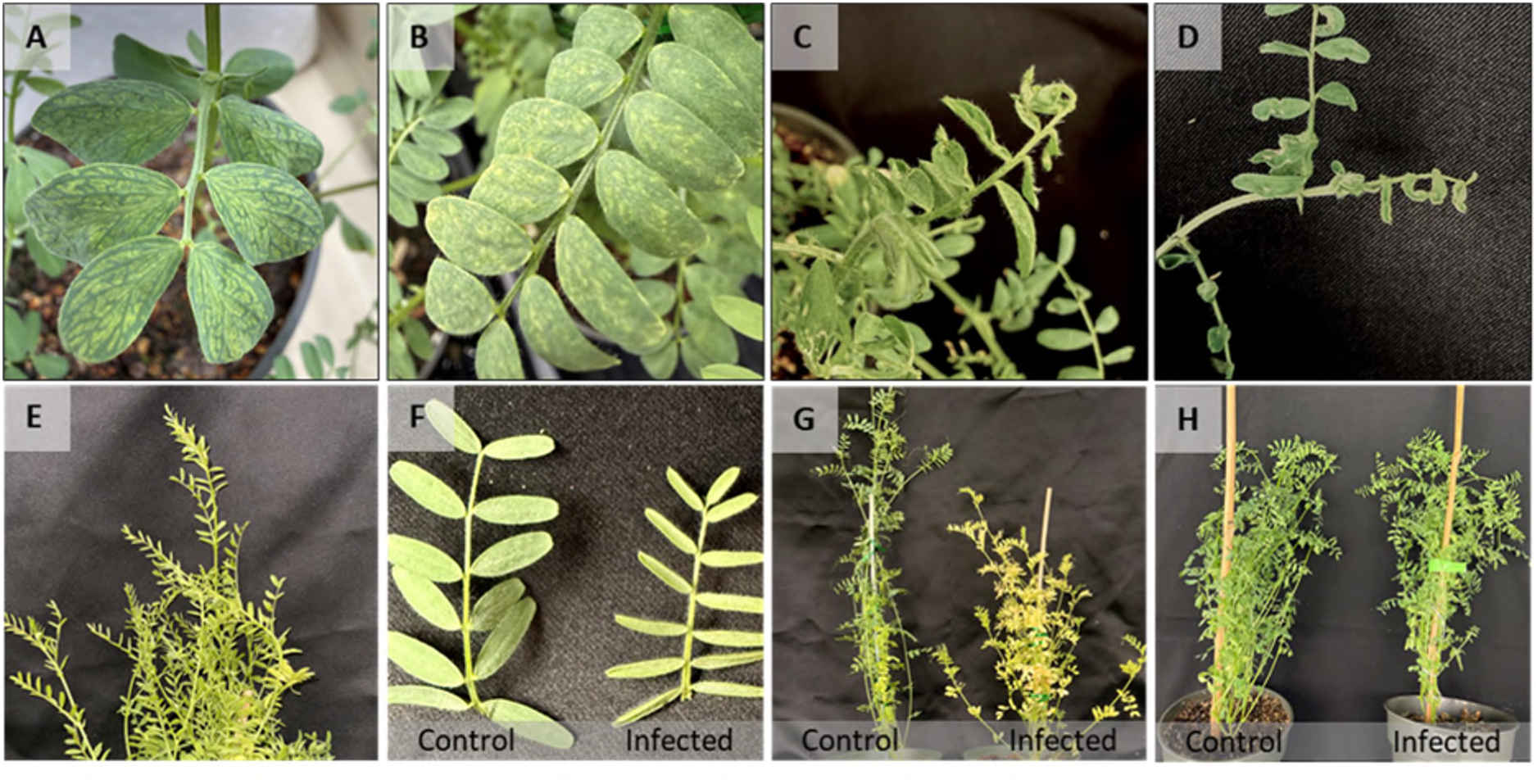
Representative symptoms observed in lentil plants 30 days after PSbMV infection. Mosaic patterns on leaflets (A, B), distortion and curling of leaflets (C, D), reduced leaflet size (E, F), dwarfing of PSbMV-inoculated plants (G) compared with mock-inoculated plants, and virus-inoculated non-symptomatic plant (H).

Out of the 86 lentil lines evaluated, six lines (CDC KR-1_AGL, PI 431893_LSP_AGL, ILL 4665_AGL, ILL 5058, PI 182217_LSP_AGL, PI 4322212_LSP_AGL) exhibited substantial variation in their qPCR results among their biological replicates and were excluded from further analysis (Supplementary file 1). The remaining 80 lines showed consistent qPCR results across replicates. Consistent with the PCR results, qPCR detected PSbMV in all inoculated leaf samples, with CT values ranging from 13 to 33 for PSbMV and from 15 to 35 for the reference gene *LceEF1α* (Supplementary file 1). Similarly, in upper systemically infected leaves, CT values ranged from 12 to 39 for PSbMV and from 14 to 28 for *LceEF1α*. Although variation in *LceEF1α* CT values was relatively small, the observed differences among samples indicate variation in cDNA input between reactions. Therefore, PSbMV transcript levels were normalized to the plant reference gene (*LceEF1α*), and ΔCT values were calculated for each line, with lower ΔCT values indicating higher virus titers and higher ΔCT values indicating lower virus titers. The 80 lentil lines exhibited a continuous distribution of ΔCT values, ranging from -14 to 12 in inoculated leaves and from -5 to 20 in systemically infected leaves (Fig. 4; Supplementary file 1). Overall, virus accumulation was high among lentil lines in the inoculated leaves, confirming successful inoculation. In contrast, a substantially wider range of virus titer was observed in the upper, systemically infected leaves. Moreover, none of the tested lentil lines exhibited complete immunity that restricts viral replication in the inoculated tissue. Rather, resistance appears to limit viral replication and/or systemic movement, resulting in markedly reduced virus accumulation in the upper leaves of some lentil lines. Since the ΔCT values did not clearly separate lines into two distinct groups, a ΔCT cutoff of 5 was arbitrarily chosen, corresponding to a high PSbMV CT value (approximately 30), to distinguish resistant and susceptible lentil lines. As expected, all previously reported resistant lentil lines (PI 212610, PI 251786, PI 297745, and PI 368648; Haddad et al. 1978) were classified as resistant.

**Fig. 4.**
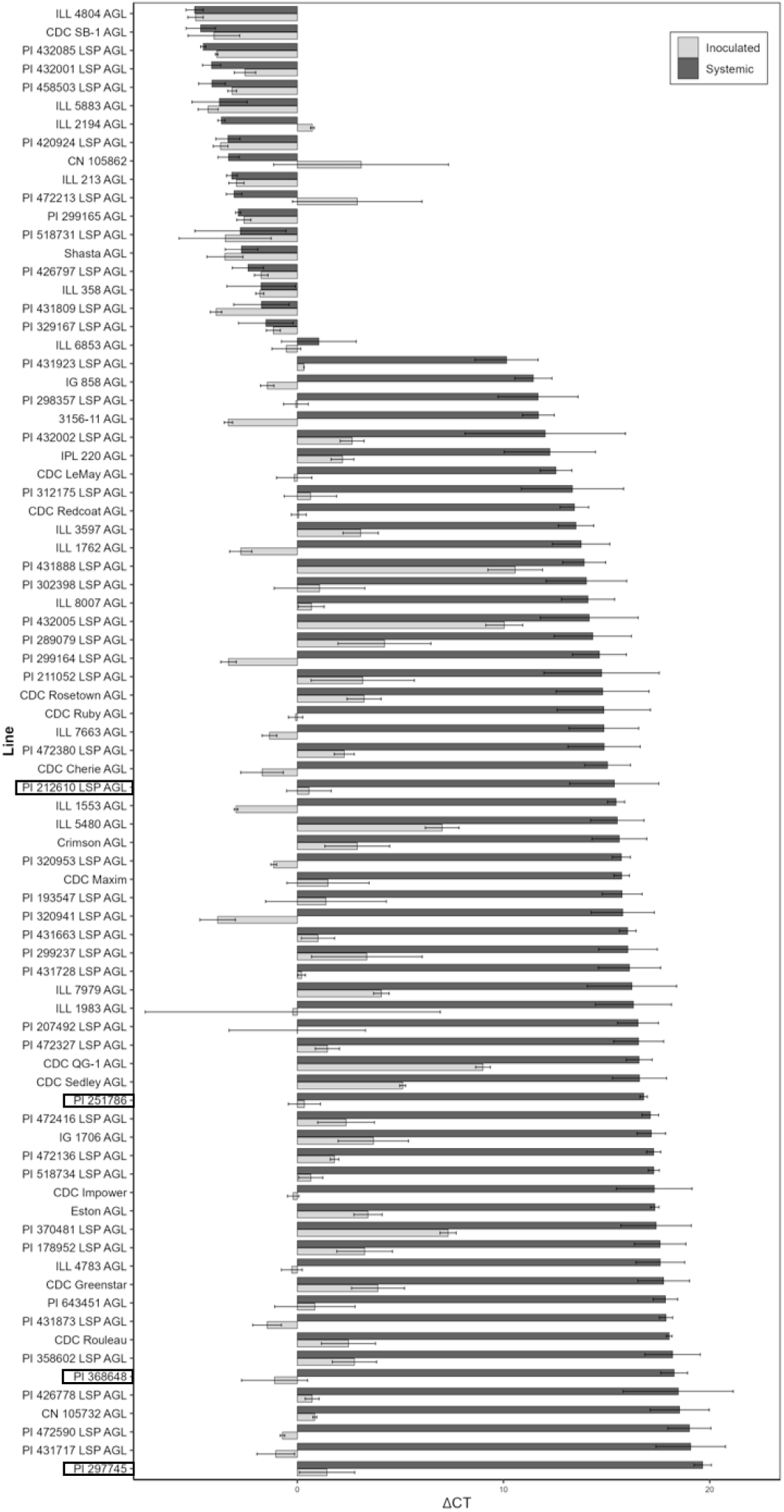
PSbMV titers across 80 lentil lines measured by qPCR. Bar plot showing mean ΔCT values for PSbMV measured in inoculated (grey bars) and systemic (black bars) leaves of lentil lines. ΔCT values were calculated as the difference between PSbMV CT and lentil reference gene (*LceEF1α*) CT. Bars represent the average ΔCT per line across three biological replicates, with higher ΔCT values indicating lower viral titer and greater resistance, and lower ΔCT values indicating higher viral titer and susceptibility. Standard errors are represented by black lines on each bar. The lines highlighted by a rectangular box are the resistant lines from the study of Haddad et al. (1978).

### *eIF4E* polymorphism in PSbMV-susceptible and resistant lentil lines resembles that reported in pea

The presence of lentil lines carrying the recessive resistance allele *sbv* within the resistant group led us to hypothesize that *eIF4E*-associated resistance contributes to the observed reduction in viral titers in the systemically infected leaves. Partial nucleotide sequences of the *eIF4E* of seven lentil lines from the resistant group (CDC Greenstar, CDC Impower, PI 518734 LSP, PI 297745, PI 368648, CDC Maxim, and CDC Crimson) and five lines from the susceptible group (CN 105862, ILL 213 AGL, ILL 2194 AGL, PI 329167 LSP AGL, and PI 472213 LSP AGL) were generated and aligned in SnapGene (v8.2.2) using the default settings of Clustal Omega, keeping the CDC Redberry *eIF4E* protein coding sequence as the reference sequence. Alignment revealed seven nucleotide polymorphisms (C64A, T189C, C221A, G319A, G322A, T336C, and G352A) that differentiated the lines of resistant group from the susceptible group (Fig. 5, Supplementary file 2). Of these, three nucleotide substitutions, C64A, T189C, and T336C, were synonymous, while the other four substitutions resulted in amino acid changes that differed between the resistant and susceptible groups. Nucleotide substitutions C221A, G319A, G322A, and G352A corresponded to amino acid substitutions A74D, G107R, A108T, and E118K, respectively, distinguishing members of the susceptible group from those of the resistant group (Fig. 6).

**Fig. 5.**
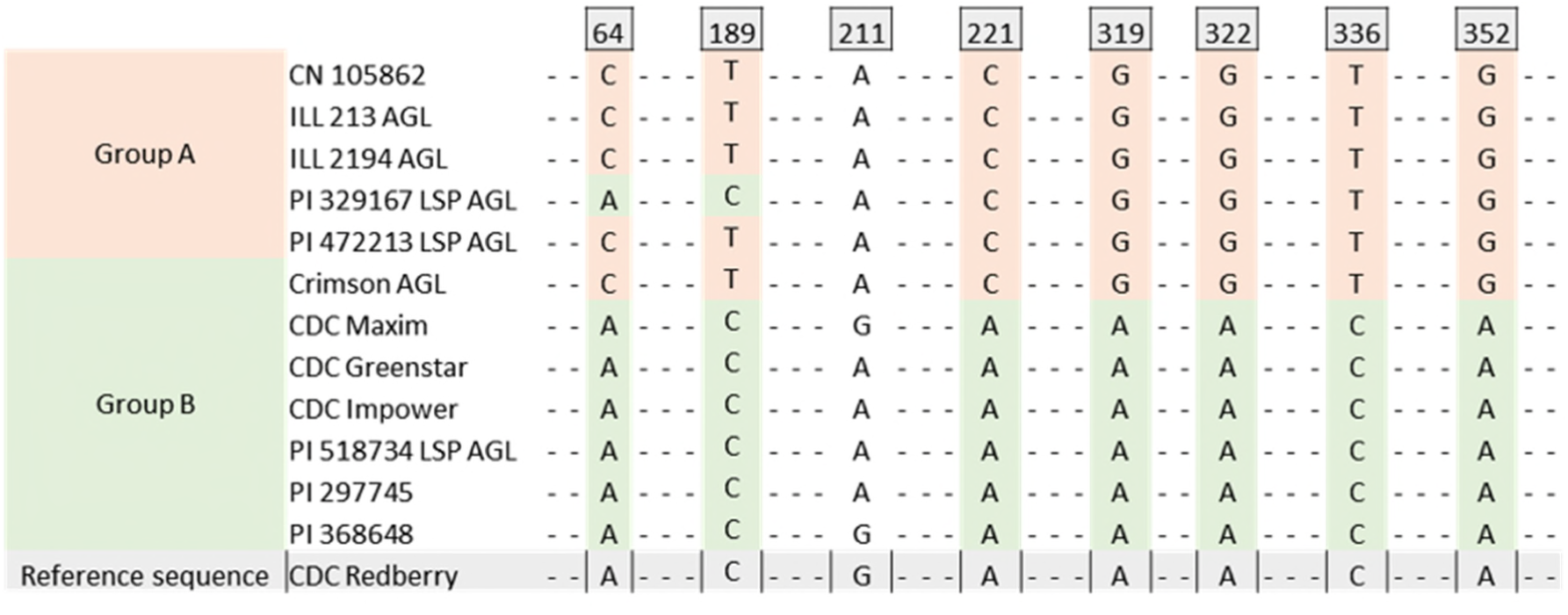
Nucleotide polymorphisms in the lentil eIF4E protein coding sequence associated with resistance to PSbMV. Nucleotides identical to those in the reference sequence are indicated by hyphens (–). Nucleotide positions are based on the CDC Redberry eIF4E coding sequence, with the adenine (A) of the start codon (ATG) designated as nucleotide position 1. Here, Group A represents susceptible lines and Group B represents resistant lines.

**Fig. 6.**
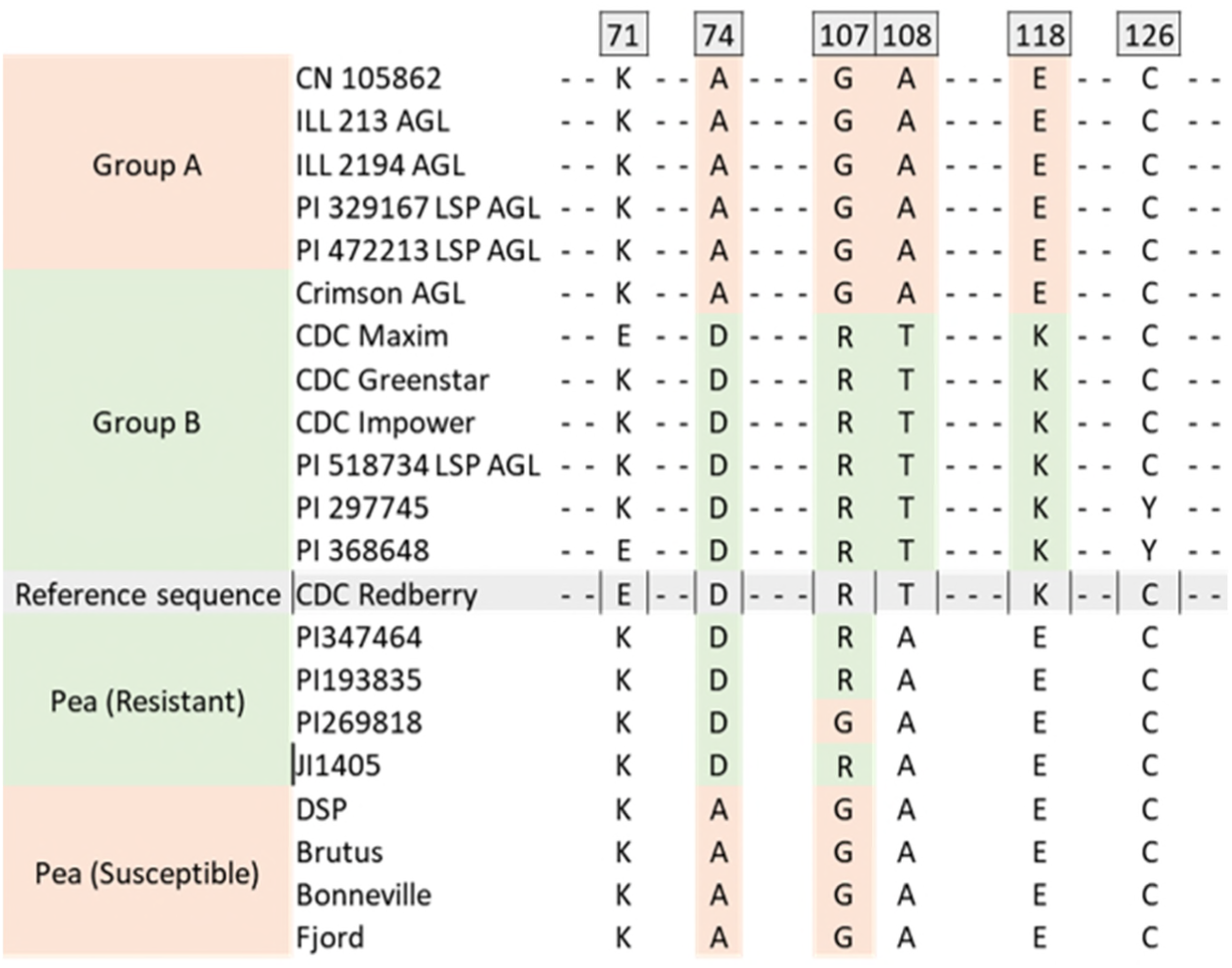
Comparison of amino acid substitutions in lentil and pea eIF4E between resistant and susceptible lines. “-” represents a similar amino acid in all sequences. Similar nucleotide polymorphisms in lentil and pea are observed at positions 221 and 319, corresponding to amino acid substitutions at positions 74 and 107, relative to the reference eIF4E sequence of the CDC Redberry lentil line. Here, Group A represents susceptible lines, and Group B represents resistant lines.

Alignment of the amino acid sequences of lentil eIF4E with the resistant and susceptible pea eIF4E sequences showed a high degree of amino acid similarity between the species. (Fig. 6, Supplementary file 2). Some of the amino acid changes associated with resistance in pea (A74D and G107R) were also found in the resistant lentil lines at their corresponding positions (Fig. 6). Protein homology modeling of lentil eIF4E from resistant and susceptible lines showed high structural similarity, including in the predicted cap-binding region (Fig. 7).

**Fig. 7.**
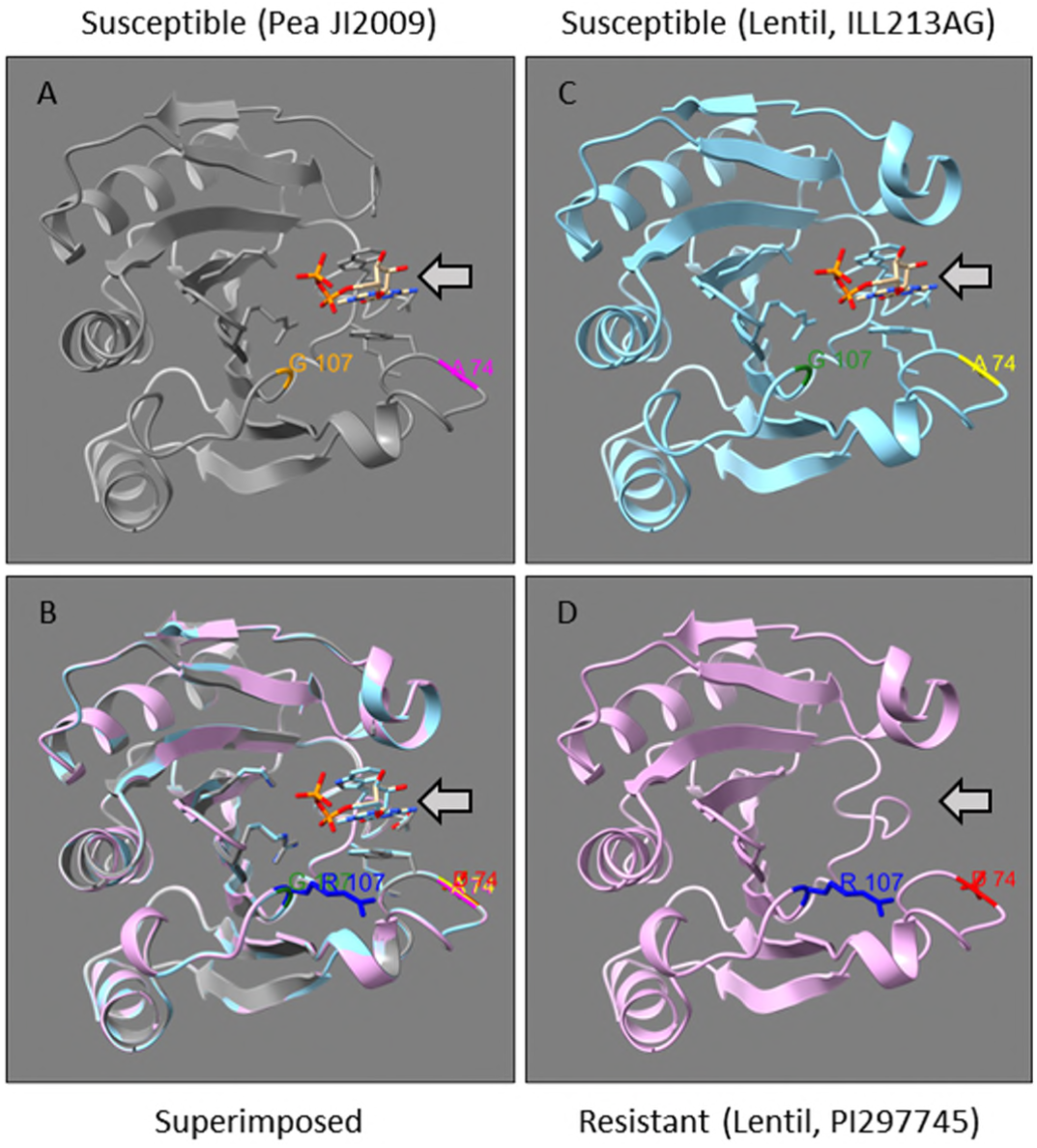
Structural modeling of eIF4E proteins from susceptible and resistant lentil lines. The eIF4E protein from the PSbMV-susceptible pea line JI2009 (A) was used as the template for homology modeling. Superimposition of the modeled proteins (B) revealed a high degree of structural similarity between pea and lentil eIF4E proteins. Amino acid substitutions associated with resistance (A74D and G107R) are highlighted in the susceptible lentil line ILL213AGL (C), the resistant lentil line PI297745 (D), and the pea template (A). Notably, the amino acid substitutions in the resistant lentil line result in the loss of the predicted binding site for the m7GTP cap analog (arrow).

### Development and validation of *eIF4E*-based KASP markers for lentil resistance genotyping

Among the four polymorphisms detected in the sequence alignment, C221A, G319A, and G352A were selected for marker development. Accordingly, three KASP markers, LceIF4E-K1, LceIF4E-K2, and LceIF4E-K3, were developed to target the corresponding loci (Table 2, Fig. 8). A total of 80 lentil lines were evaluated using three KASP markers and the clustering pattern clearly separated the lines into two groups corresponding to C or A, G or A, and G or A polymorphisms targeted by the LceIF4E-K1, LceIF4E-K2, LceIF4E-K3 markers at the C221A, G319A, and G352A nucleotide positions, respectively (Fig. 9). Furthermore, these allele discriminations were consistent with the expected results based on the *eIF4E* nucleotide sequences of the 12 individuals sequenced in this study (Fig. 8).

**Fig. 8.**
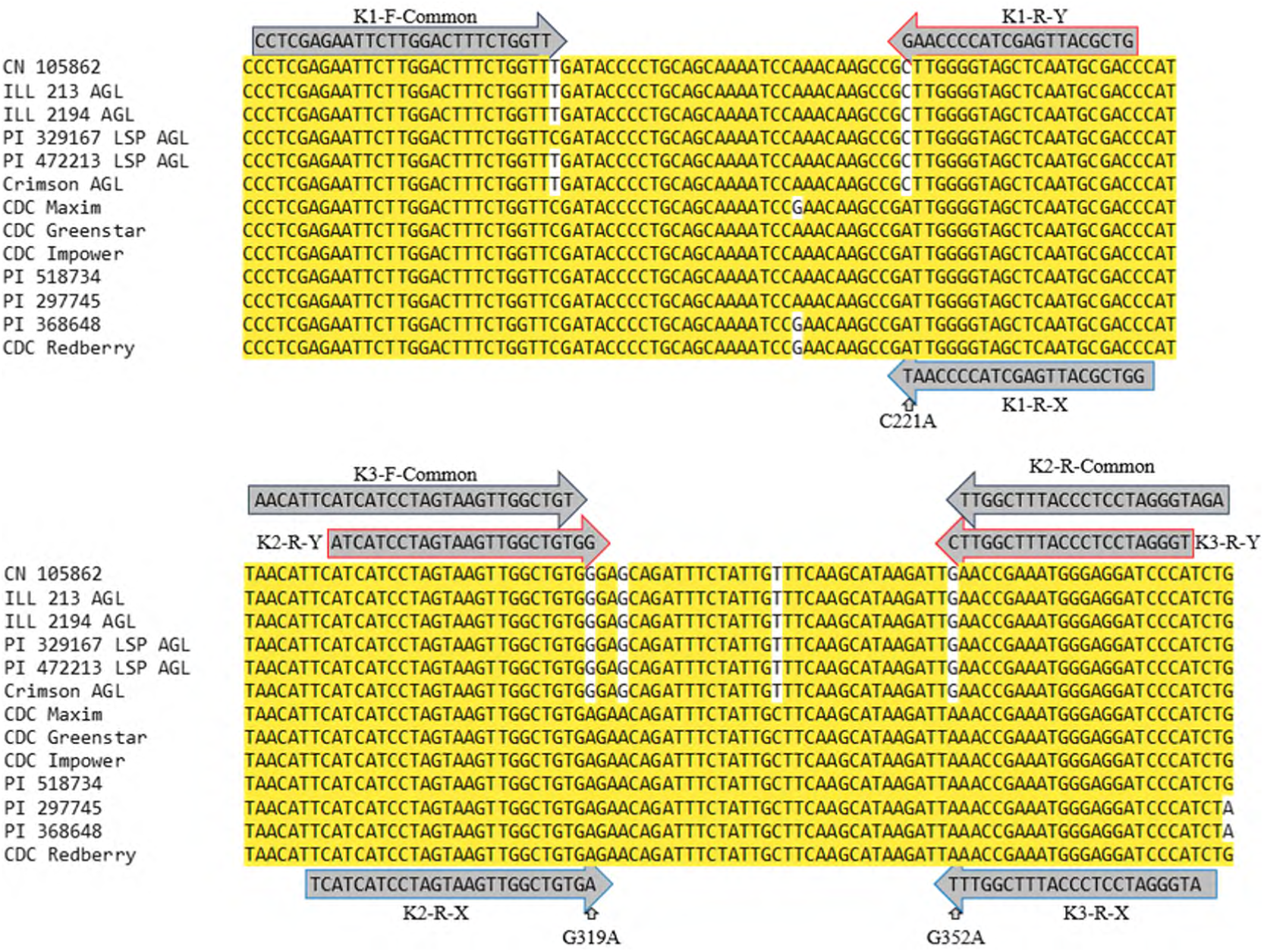
Position and sequence context of the developed KASP marker within aligned *eIF4E* sequences of susceptible and resistant lentil lines. The polymorphic site used for marker design is indicated relative to its coordinate in the CDC Redberry eIF4E reference sequence.

**Fig. 9.**
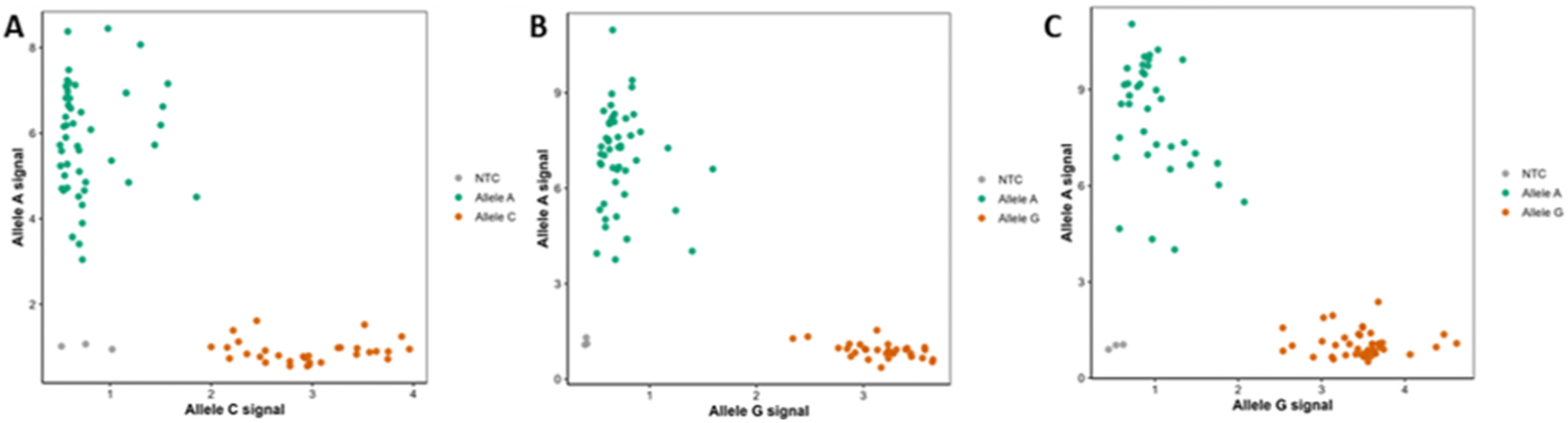
Cluster plot showing allele discrimination of 80 lentil lines genotyped using KASP markers: LceIF4E-KASP1 (A), LceIF4E-K2 (B). LceIF4E-K3 (C). The fluorescence intensity of the FAM probe (allele “A”, resistant) is plotted on the y-axis, and the HEX probe (allele “G” or “C”, susceptible) on the x-axis. Each point represents an individual genotype. Samples cluster into two distinct groups corresponding to homozygous resistant (A/A; green) and homozygous susceptible (G/G or C/C; red) genotypes. Water/Negative controls (NC) are shown in grey dots.

Of the 86 lines, six lines (CDC KR-1 AGL, PI 431893 LSP AGL, ILL 4665 AGL, ILL 5058, PI 182217 LSP AGL, PI 4322212 LSP AGL) that showed inconsistent qPCR results also showed similar inconsistent allele calls in at least one replicate (Supplementary file 1). These inconsistencies suggest a heterogeneous seed mixture, containing seeds with both homozygous susceptible and resistant eIF4E alleles. It is not uncommon to find heterogeneous seed mixtures in some lines. To confirm the presence of this heterogeneity, Sanger sequencing on cDNA from three replicates was performed using LceIF4E-specific primers described above. As expected, a single-nucleotide call at these positions matched the results from the KASP marker and qPCR (Fig. 10, Supplementary file 2), confirming the seed heterogeneity in some lines.

**Fig. 10.**
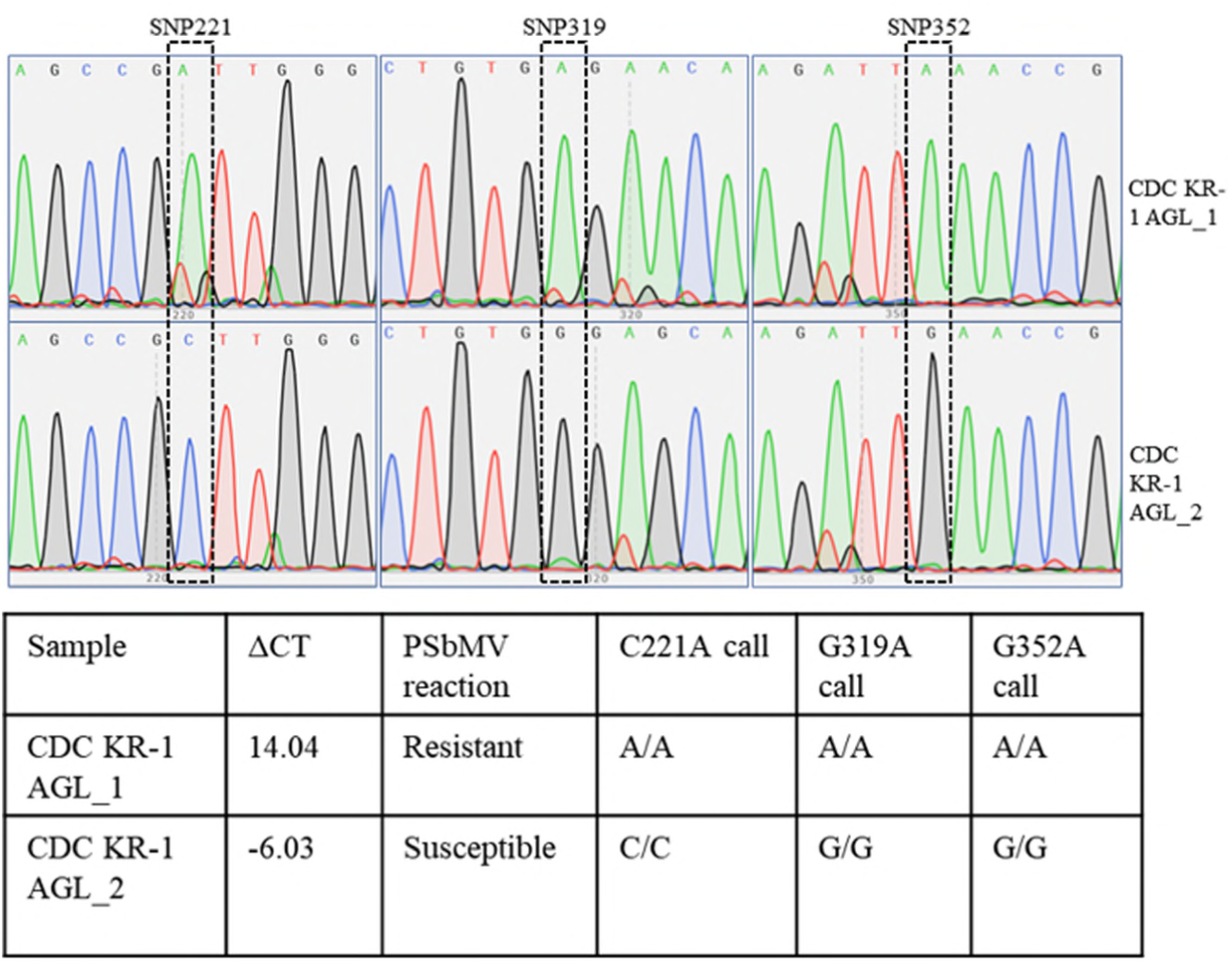
Chromatogram of Sanger sequencing results of the eIF4E gene region for a representative lentil line (CDC KR-1 AGL) showing inconsistent genotype calls in KASP analysis. The table below shows the qPCR ΔCT value, reaction to PSbMV, and KASP allele calls of those two samples.

The remaining 80 lentil lines were successfully genotyped using the KASP assay, which yielded consistent genotype calls across three biological replicates (Tables 3,4,5). Out of the total 80, 19 qPCR-determined susceptible lines consistently showed susceptible alleles (C/C, G/G, and G/G) at all three tested nucleotide positions (221, 319, and 352) (Table 3, Supplementary file 1). Interestingly, six qPCR-determined resistant lines, including the eIF4E Sanger-sequenced Crimson AGL, showed susceptibility-associated alleles at all three KASP-assessed positions (Table 4), suggesting they carry an additional mechanism of resistance.

**Table 3:**
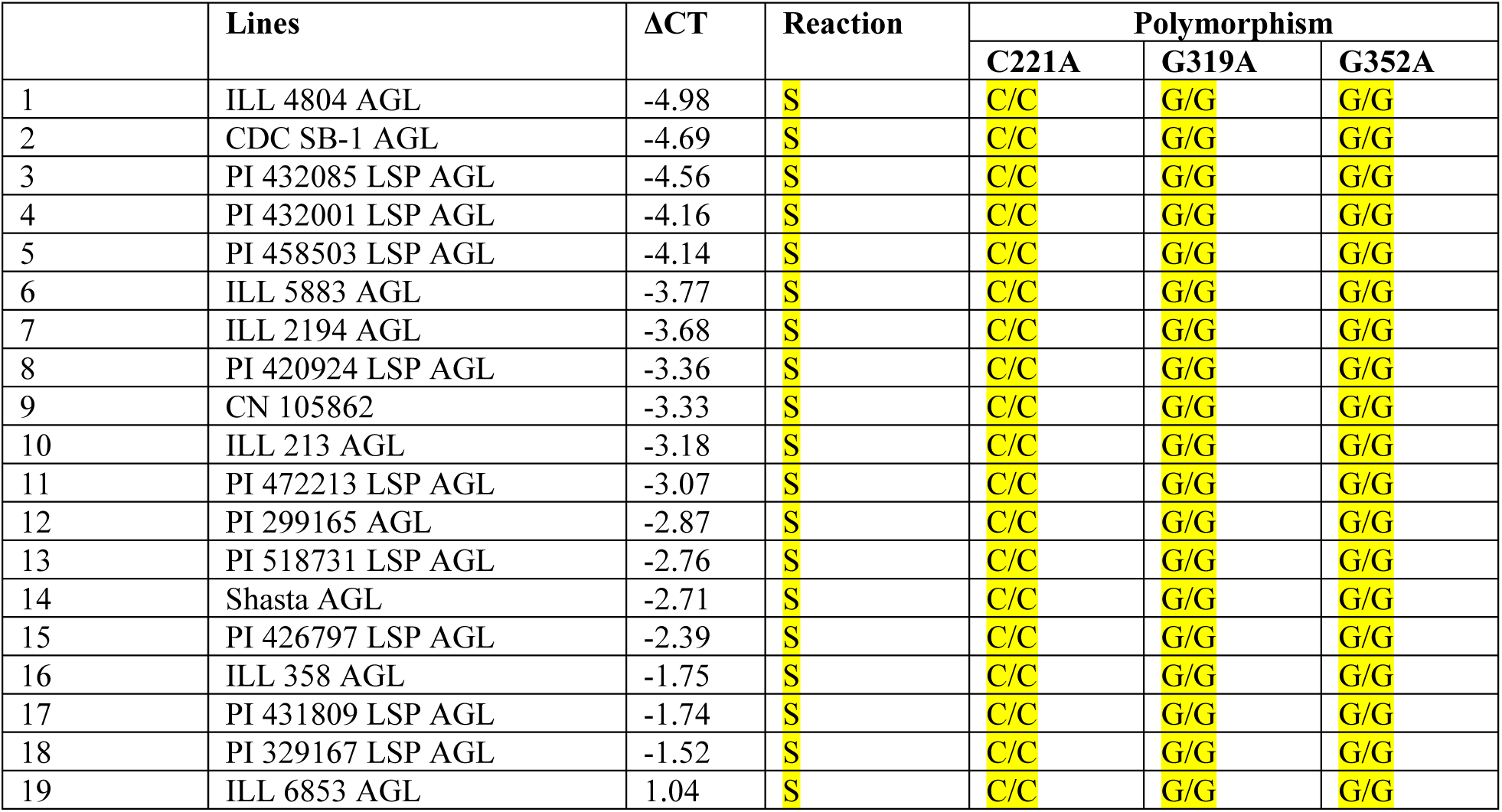
Genotype calls of lines showing a susceptible reaction to PSbMV for single nucleotide polymorphisms C221A, G319A, and G352A, identified with the KASP markers LceIF4E-K1, LceIF4E-K2, and LceIF4E-K3, respectively, and their qPCR ΔCT values. Yellow color coding represents the PSbMV-susceptible allele, while grey represents the resistant allele.

**Table 4:** Lines showing resistant reaction to PSbMV but having non-resistant alleles at nucleotide positions 221, 319, and 352 identified by KASP markers LceIF4E-K1, LceIF4E-K2, and LceIF4E-K3, respectively. Yellow color coding represents the PSbMV-susceptible allele, while grey represents the resistant allele.

| | Lines | ΔCT | Reaction | Polymorphism | | |
| --- | --- | --- | --- | --- | --- | --- |
|  |  |  |  | C221A | G319A | G352A |
| 1 | PI 302398 LSP AGL | 14.02 | R | C/C | G/G | G/G |
| 2 | PI 299164 LSP AGL | 14.65 | R | C/C | G/G | G/G |
| 3 | ILL 8007 AGL | 14.10 | R | C/C | G/G | G/G |
| 4 | PI 299237 LSP AGL | 16.04 | R | C/C | G/G | G/G |
| 5 | Crimson AGL | 15.62 | R | C/C | G/G | G/G |
| 6 | IG 1706 AGL | 17.17 | R | C/C | G/G | G/G |

Of the 55 qPCR-determined resistant lines, 49, 49, and 37 were identified as carrying resistance-associated alleles by the LceIF4E-K1, LceIF4E-K2, and LceIF4E-K3 markers, respectively, with 30 lines possessing resistance-associated alleles at all three marker loci and remaining lines having at least one resistance-associated allele in any of the other two loci (Table 5, Supplementary file 1). The presence of a resistance-associated allele detected by any one of these markers was associated with resistance. Furthermore, the combined presence of resistance-associated alleles detected by LceIF4E-K1 and LceIF4E-K2 was sufficient to predict resistance, even in the absence of the LceIF4E-K3 resistance-associated allele.

**Table 5:**
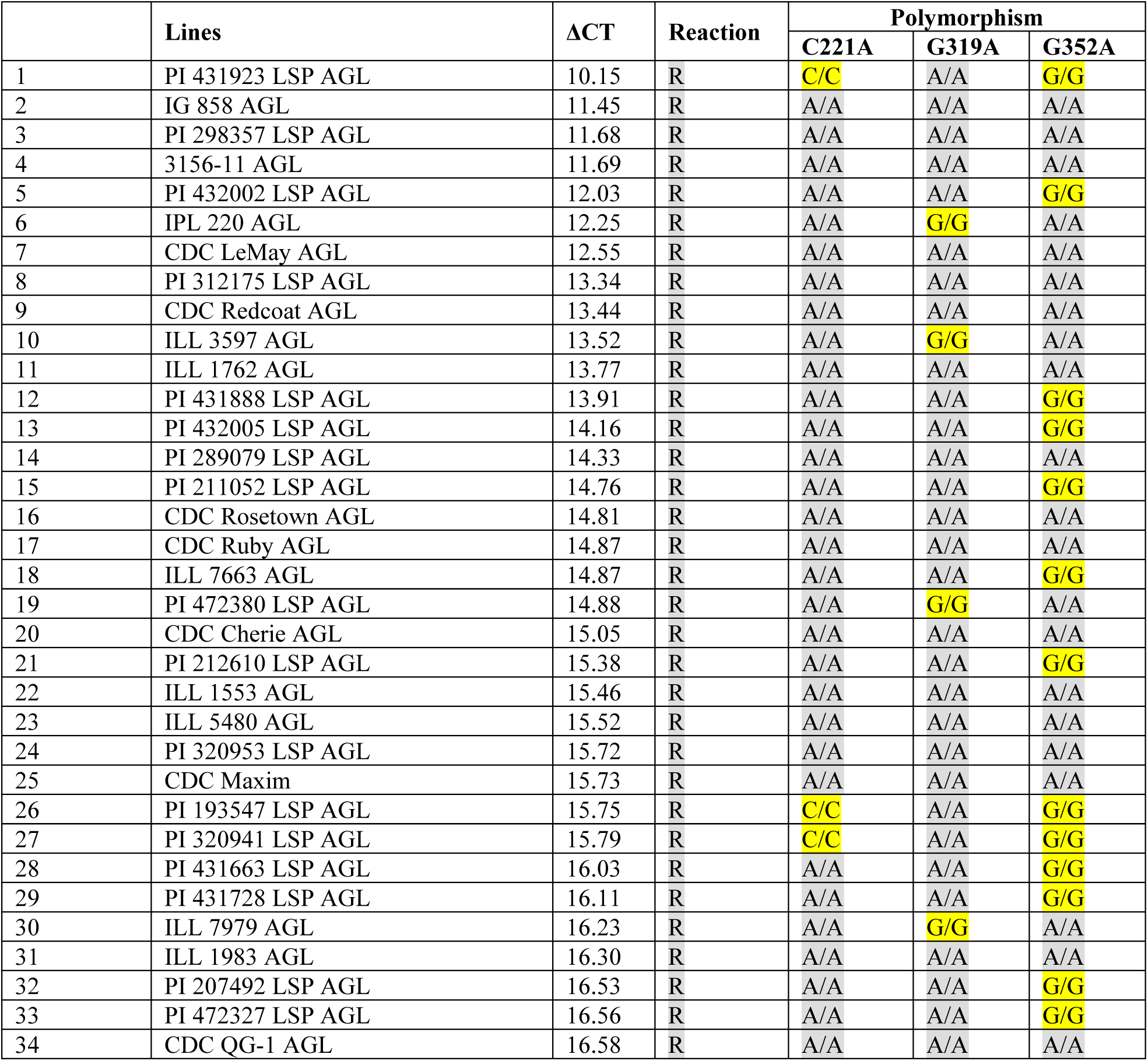
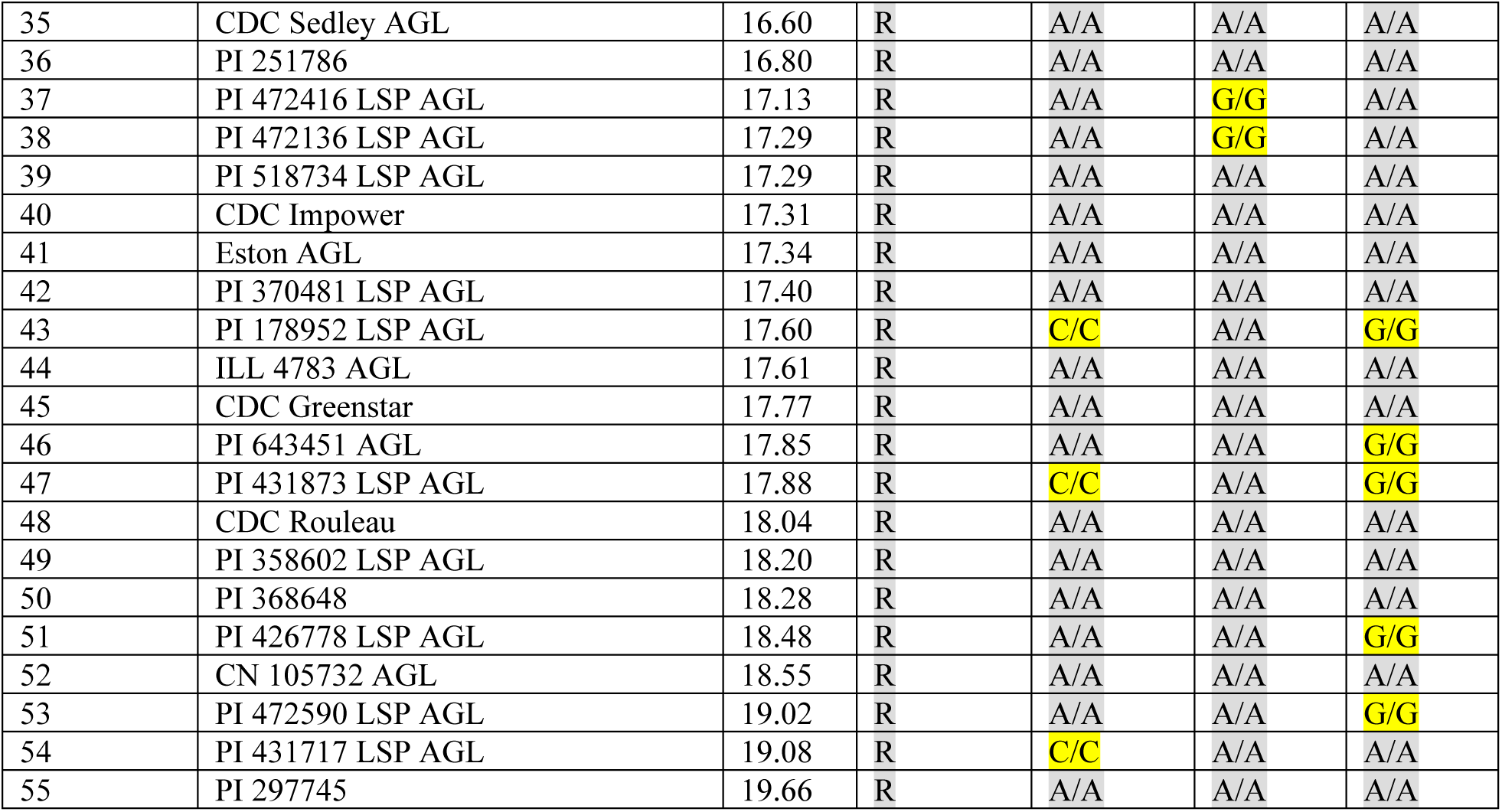
Genotype calls of PSbMV-resistant lentil lines, their qPCR ΔCT values, and polymorphisms at nucleotide positions 221, 319, and 352, identified by the KASP markers LceIF4E-K1, LceIF4E-K2, and LceIF4E-K3, respectively. Yellow color coding represents the PSbMV-susceptible allele, while grey represents the resistant allele.

### Validation of KASP markers on the genomic DNA in lentil

Often, KASP assays are performed on genomic DNA, whereas assays in this study were performed on intron-free cDNA. Introns in genomic DNA, especially if they are present between the KASP primers, could affect the usability of KASP markers designed from cDNA. To verify if these KASP markers can be used with genomic DNA, the genomic sequence of *eIF4E* from reference line (CDC Redberry) was examined and four introns were identified, but none were located within the amplicon targeted by the forward and reverse KASP primers (Fig. 11). A subset of 25 lines was randomly selected from the 80 lines (Table 6), and KASP genotyping was conducted using three KASP markers following the same protocol described above for the cDNA. Genotype calls from gDNA showed results similar to those previously generated from cDNA (Fig. 12, Table 6), consistent with the cDNA-based results above.

**Fig. 11.**
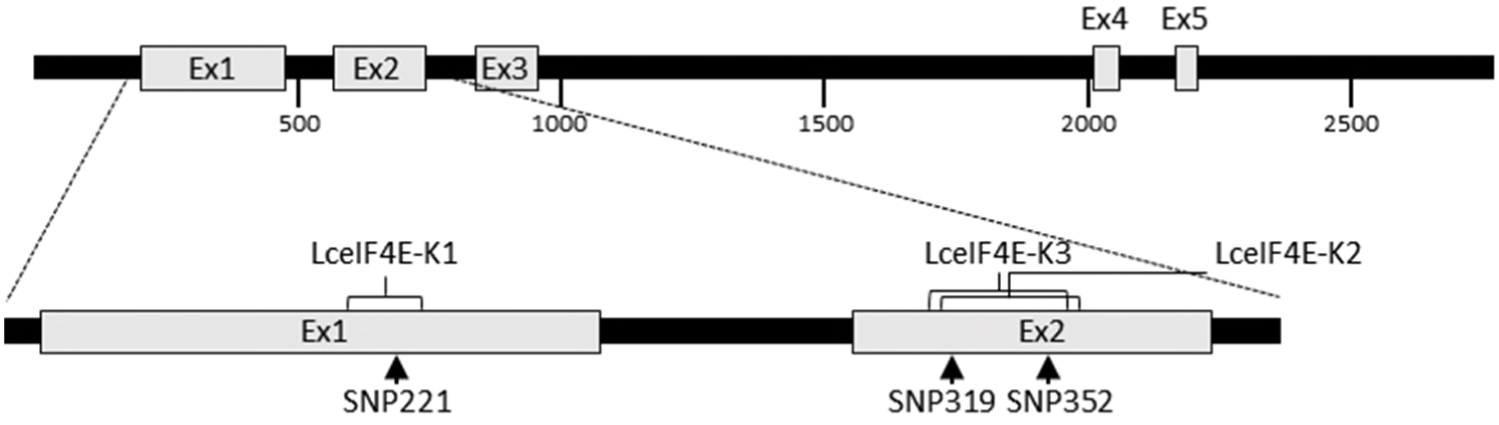
Schematic representation of the lentil eIF4E gene structure and the positions of the KASP marker amplicons. Exons (Ex) are shown as boxes and introns as connecting black lines. Lines inside exons represent the SNPs position and brackets at the top of exons labeled with the respective KASP marker names indicate the amplified regions.

**Fig. 12.**
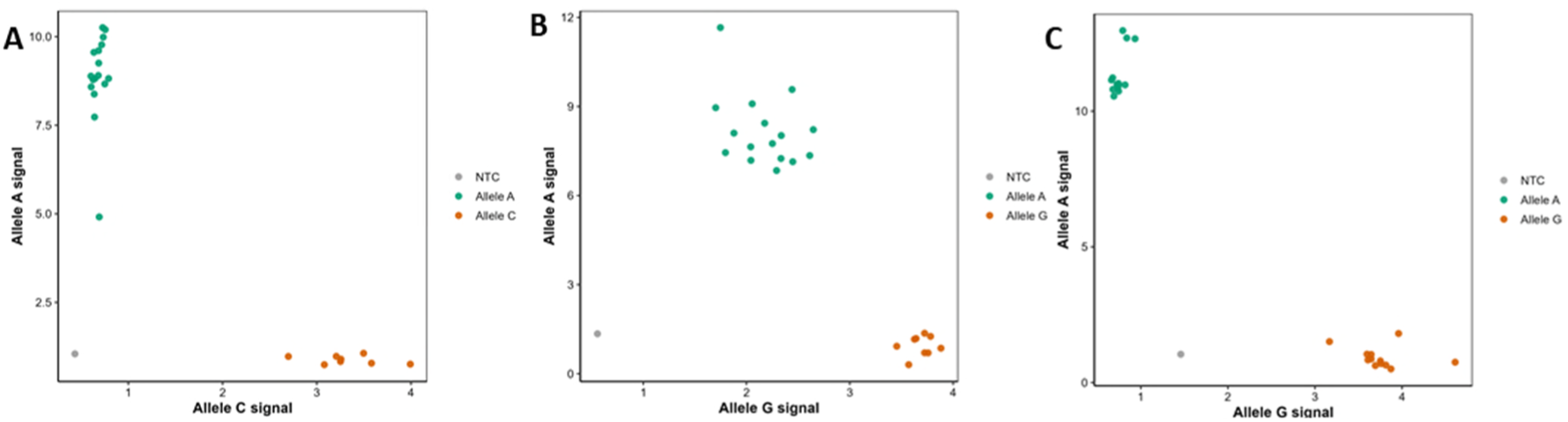
KASP cluster plot showing allele discrimination of 25 lentil line genotypes using KASP markers; LceIF4E-KASP1 (A), LceIF4E-K2 (B). LceIF4E-K3 (C). Fluorescence intensity of the FAM probe (allele “A”, resistant) is plotted on the y-axis, and HEX probe (allele “G” or “C”, susceptible) on the x-axis. Each point represents an individual genotype. Samples cluster into two distinct groups corresponding to homozygous resistant (A/A; green) and homozygous susceptible (G/G or C/C; red) genotypes. Water/Negative controls (NC) are shown in grey dots.

**Table 6:**
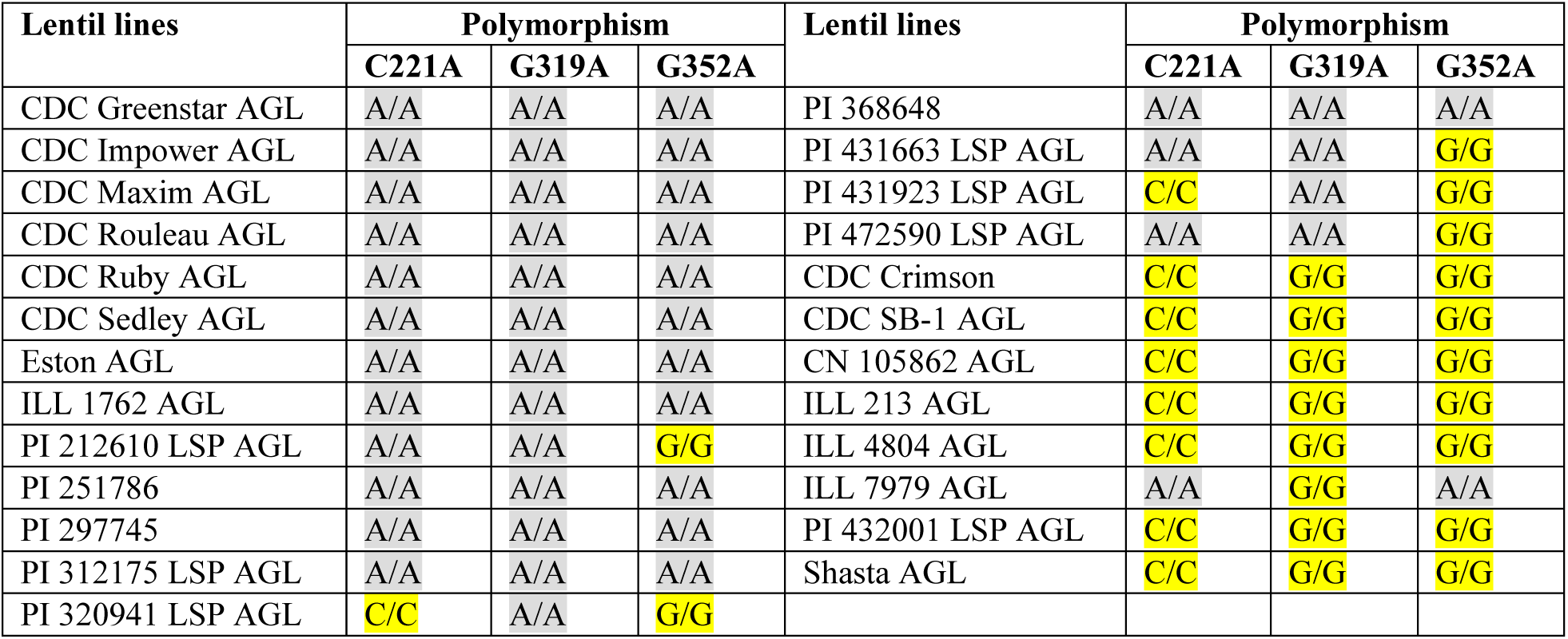
Genotype calls for 25 randomly selected lines using genomic DNA, targeting single-nucleotide polymorphisms observed at C221A, G319A, and G352A with KASP markers LceIF4E-K1, K-2, and K3, respectively. Yellow color coding represents the PSbMV-susceptible allele, while grey represents the resistant allele.

| Lentil lines | Polymorphism |  |  | Lentil lines | Polymorphism |  |  |
| --- | --- | --- | --- | --- | --- | --- | --- |
|  | C221A | G319A | G352A |  | C221A | G319A | G352A |
| CDC Greenstar AGL | A/A | A/A | A/A | PI 368648 | A/A | A/A | A/A |
| CDC Impower AGL | A/A | A/A | A/A | PI 431663 LSP AGL | A/A | A/A | G/G |
| CDC Maxim AGL | A/A | A/A | A/A | PI 431923 LSP AGL | C/C | A/A | G/G |
| CDC Rouleau AGL | A/A | A/A | A/A | PI 472590 LSP AGL | A/A | A/A | G/G |
| CDC Ruby AGL | A/A | A/A | A/A | CDC Crimson | C/C | G/G | G/G |
| CDC Sedley AGL | A/A | A/A | A/A | CDC SB-1 AGL | C/C | G/G | G/G |
| Eston AGL | A/A | A/A | A/A | CN 105862 AGL | C/C | G/G | G/G |
| ILL 1762 AGL | A/A | A/A | A/A | ILL 213 AGL | C/C | G/G | G/G |
| PI 212610 LSP AGL | A/A | A/A | G/G | ILL 4804 AGL | C/C | G/G | G/G |
| PI 251786 | A/A | A/A | A/A | ILL 7979 AGL | A/A | G/G | A/A |
| PI 297745 | A/A | A/A | A/A | PI 432001 LSP AGL | C/C | G/G | G/G |
| PI 312175 LSP AGL | A/A | A/A | A/A | Shasta AGL | C/C | G/G | G/G |
| PI 320941 LSP AGL | C/C | A/A | G/G |  |  |  |  |

## Discussion

### qPCR-based screening of PSbMV resistance in lentil

In the search for naturally occurring resistance, a lentil diversity panel was screened for resistance to PSbMV using rub inoculation experiments in growth chambers. Symptomatic PSbMV-inoculated lentil lines exhibited milder symptoms, making it difficult to visually score the resistance. However, when virus presence was assessed in inoculated leaves at 30 days post-inoculation (dpi) by both conventional PCR and qPCR, all samples tested positive (Fig. 4), suggesting that the lentil lines are permissive to PSbMV replication and movement within the inoculated leaves. All upper, systemically infected leaves of PSbMV-inoculated lentil plants tested positive for PSbMV by qPCR. Nevertheless, some lines exhibited markedly lower virus titers than others, indicating that resistance may act by restricting viral replication, limiting systemic movement, promoting viral RNA degradation, or through a combination of these mechanisms. Given the presence of previously reported recessive-resistance lentil lines in this resistance group and a similar resistance mechanism reported in pea, it was hypothesized that recessive resistance might explain the low virus titers observed in these lines.

The screening detected PSbMV in both inoculated and systemically infected leaves, including in the systemically infected leaves of resistant lines, though at a lower titer. This observation differs from previous reports of recessive resistance in pea and lentil, where resistance, as assessed by visual symptom evaluation or ELISA screening, was associated with the absence of PSbMV (Haddad et al. 1978; Konečná et al. 2014; Swisher Grimm and Porter 2020, 2021). It is suspected that these previous observations resulted from low virus titers in resistant plants rather than the complete absence of qPCR-detectable viral nucleic acids. In this study, PSbMV-specific probes that had previously been validated to detect PSbMV even after a 10⁻⁶ dilution were used (Paudel et al. 2025). This level of sensitivity is likely greater than that of ELISA-based assays, which are generally regarded as less sensitive than qPCR for virus detection (Bertolini et al. 2008; Gachon et al. 2004; Torre et al. 2020). The presence of low levels of virus in the resistant lines is also supported by low but detectable GUS expression from the chimeric PSbMV construct in the resistance line (Gao et al. 2004a). Similarly, low detectable potyvirus (tobacco etch virus) was also shown in pepper plants possessing *eIF4E*-associated recessive resistance (Kang et al. 2005). Together, the presence of lower but qPCR-detectable potyviruses in *eIF4E*-associated resistant lines is further supported by these observations and the existing literature. Furthermore, these assays were performed on systemically infected leaves, located above the inoculated leaves at 14 days post-inoculation. Because reduced viral replication and systemic movement in the resistant lines are indicated by these results, viral titers are expected to decline further in the upper leaves.

### *eIF4E*-associated resistance in lentils

Recessive resistance results from mutations in host susceptibility genes required for virus replication, translation, and movement (Nicaise 2014; Sanfaçon 2015). Recessive resistance against potyviruses is mostly associated with the translation initiation factors, particularly eIF4E or its isoform eIF4E(iso)4E (Zlobin and Taranov 2023). Consistent with this, the only characterized recessive resistance gene (*sbm*) against PSbMV in pea is associated with changes in the amino acid sequence of the eIF4E protein (Konečná et al. 2014; Swisher Grimm and Porter 2020). The eukaryotic translation initiation factor eIF4E is a multifunctional host protein that, amongst other functions, interacts with the 5’ cap structure of the eukaryotic mRNA and initiates cellular mRNA translation (Mars et al. 2024; Zlobin and Taranov 2023). The potyviruses lack the 5’ cap structure but possess a viral genome-linked protein (VPg), which is known to interact with the host eIF4E and is suggested to play a similar role to that of the eukaryotic cap (Mars et al. 2024; Pollari et al. 2024; Zlobin and Taranov 2023). In addition, eIF4E is suggested to play a role in viral replication and cell-to-cell movement (Ashby et al. 2011; Gao et al. 2004b).

Plants possess at least two isoforms of the eukaryotic translation initiation factor 4E, namely eIF4E and eIF(iso)4E. These isoforms play largely redundant roles in eukaryotic mRNA translation. However, plant viruses have evolved to specifically exploit one or the other isoforms, and mutations in either eIF4E or eIF(iso)4E can confer recessive resistance to the group of viruses that depend on the corresponding isoform for successful infection (Kobayashi et al. 2014; Sanfaçon 2015; Zlobin and Taranov 2023).

Comparison of eIF4E sequences from lentil lines showed high nucleotide and protein sequence similarity to that of pea. Their predicted protein structures show a high degree of similarity, suggesting an interaction between the PSbMV VPg and the lentil eIF4E similar to that observed in pea. Furthermore, six amino acid substitutions in the resistant lentil lines were revealed by comparative analysis of eIF4E between resistant and susceptible lentil lines, of which two, A74D and G107R (Figs. 5 and 6), are also implicated in PSbMV resistance in pea (Gao et al. 2004a; Gao et al. 2004b). The G107R substitution is located near the cap-binding pocket of eIF4E and is suggested to disrupt the interaction between eIF4E and the potyvirus VPg protein, thereby conferring resistance (Kang et al. 2005; Yeam et al. 2007). In fact, amino acid substitution at this site (G107R) has been reported in recessive resistance to potyviruses in several crops (Dinkova et al. 2016). In addition to the naturally occurring amino acid substitutions, mutational analysis of pea identified a number of amino acid substitutions that result in resistance to PSbMV. However, some amino acid substitutions can reduce host translation efficiency (Ashby et al. 2011), thereby impairing host fitness and likely imposing selective pressure that favors substitutions with minimal adverse effects on the host, which are likely the amino acid substitutions commonly observed in naturally occurring resistance.

### KASP markers for the screening of resistance lines

Given the suspected presence of eIF4E-associated recessive resistance in the lentil lines used in this study, three KASP markers targeting the C221A, G319A, and G352A nucleotide changes, corresponding to the A74D, G107R, and E118K amino acid substitutions identified in resistant lines, were developed and validated. Among the three markers, LceIF4E-K1, followed by LceIF4E-K2, predicted a higher number of resistant lines compared to the third LceIF4E-K3 marker, and the combined use of two markers (LceIF4E-K1, LceIF4E-K2) predicted 100% of the resistant lines. Furthermore, the detection of heterogeneous seed mixtures across several lentil lines, their accurate genotypic classification using KASP markers, and the corresponding agreement with qPCR-based resistance phenotypes provide additional validation of the marker assays.

Interestingly, discrepancies were observed in six lines identified as susceptible by all three KASP assays but resistant in the qPCR assays. Also, higher viral titers (ΔCT ≤ 5) were detected by qPCR in all susceptible lines (19 out of 19) that carried KASP-identified susceptibility-associated alleles at all three loci. In contrast, six resistant lines which accumulated relatively low virus titers were found to have KASP-identified susceptible-associated alleles. These discrepancies could be explained by two potential mechanisms. First, these lines might have additional amino acid substitutions in eIF4E at positions not covered by the KASP markers, which could confer resistance. Indeed, mutational analyses of pea eIF4E have shown that several additional amino acid changes can confer recessive resistance to PSbMV (Ashby et al. 2011). However, the full *eIF4E* sequence generated in this study from one such line, Crimson AGL (Fig. 6, Supplementary file 2), had a susceptible *eIF4E* allele, leading to second possibility of the presence of additional resistance genes independent of *eIF4E*. Similar observations have also been reported in pea, where several lines possessing susceptible-like eIF4E amino acid sequences nevertheless exhibited resistance to PSbMV (Konečná et al. 2014).

The KASP markers developed in this study provide a rapid, reproducible, and cost-effective tool for marker-assisted selection of PSbMV resistance in lentil, akin to the previously successful use of *eIF4E*-based KASP markers in other potyvirus pathosystems (Hart and Griffiths 2013; Swisher Grimm and Porter 2020, 2021). Although, the PSbMV isolate used in this study originated from infected pea seeds collected in Saskatchewan and exhibits high nucleotide and amino acid sequence similarity to the P4 pathotype (Paudel et al. 2025), further research is needed to determine whether the amino acid substitutions identified in this study also confer resistance to other PSbMV pathotypes and additional potyviruses. Nevertheless, because the P4 pathotype is the predominant PSbMV pathotype reported in major legume-growing regions, the KASP markers developed here should provide a valuable tool for marker-assisted selection and accelerate the development of PSbMV-resistant lentil cultivars.

## Acknowledgment

Financial support for this research was provided by the Pulse Science Cluster under the Sustainable Canadian Agricultural Partnership. GRP also acknowledges the Western Grains Research Foundation for graduate scholarship support received during this research.

