## Supplementary file 1 for "Screening for Pea Seed-borne Mosaic Virus Resistance (PSbMV) in Lentil and Development of Associated KASP Markers"

Lc_S_CN 105862	1	------------------CCTAAATCCATCATCACCGACGATCAAATCACAGCAAACCCTAATCGGGTTATCGAAGACGACAACAATCTTGAAGAAGGAGAGATCCTCGATGAAGACGATTCCTCCGCCA
Lc_S_ILL 213AGL	1	ATGGTTGTAGAAGACACCCCTAAATCCATCATCACCGACGATCAAATCACAGCAAACCCTAATCGGGTTATCGAAGACGACAACAATCTTGAAGAAGGAGAGATCCTCGATGAAGACGATTCCTCCGCCA
Lc_S_ILL 2194	1	-----------------CCCTAAATCCATCATCACCGACGATCAAATCACAGCAAACCCTAATCGGGTTATCGAAGACGACAACAATCTTGAAGAAGGAGAGATCCTCGATGAAGACGATTCCTCCGCCA
Lc_S_PI 329167	1	ATGGTTGTAGAAGACACCCCTAAATCCATCATCACCGACGATCAAATCACAGCAAACCCTAATAGGGTTATCGAAGACGACAACAATCTTGAAGAAGGAGAGATCCTCGATGAAGACGATTCCTCCGCCA
Lc_S_PI 472213	1	ATGGTTGTAGAAGACACCCCTAAATCCATCATCACCGACGATCAAATCACAGCAAACCCTAATCGGGTTATCGAAGACGACAACAATCTTGAAGAAGGAGAGATCCTCGATGAAGACGATTCCTCCGCCA
Lc_R_CDC Crimson	1	-------------------CTAAATCCATCATCACCGACGATCAAATCACAGCAAACCCTAATCGGGTTATCGAAGACGACAACAATCTTGAAGAAGGAGAGATCCTCGATGAAGACGATTCCTCCGCCA
Lc_R_CDC Maxim	1	--------------------TAAATCCATCATCACCGACGATCAAATCACAGCAAACCCTAATAGGGTTATCGAAGACGACAACAATCTTGAAGAAGGAGAGATCCTCGATGAAGACGATTCCTCCGCCA
Lc_R_CDC Greenstar 1 ATGGTTGTAGAAGACACCCCTAAATCCATCATCACCGACGATCAAATCACAGCAAACCCTAATAGGGTTATCGAAGACGACAACAATCTTGAAGAAGGAGAGATCCTCGATGAAGACGATTCCTCCGCCA
Lc_R_CDC Impower	1	----------------------AATCCATCATCACCGACGATCAAATCACAGCAAACCCTAATAGGGTTATCGAAGACGACAACAATCTTGAAGAAGGAGAGATCCTCGATGAAGACGATTCCTCCGCCA
Lc_R_PI 518734	1	ATGGTTGTAGAAGACACCCCTAAATCCATCATCACCGACGATCAAATCACAGCAAACCCTAATAGGGTTATCGAAGACGACAACAATCTTGAAGAAGGAGAGATCCTCGATGAAGACGATTCCTCCGCCA
Lc_R_PI 297745	1	ATGGTTGTAGAAGACACCCCTAAATCCATCATCACCGACGATCAAATCACAGCAAACCCTAATAGGGTTATCGAAGACGACAACAATCTTGAAGAAGGAGAGATCCTCGATGAAGACGATTCCTCCGCCA
Lc_R_PI 368648	1	-----------------CCCTAAATCCATCATCACCGACGATCAAATCACAGCAAACCCTAATAGGGTTATCGAAGACGACAACAATCTTGAAGAAGGAGAGATCCTCGATGAAGACGATTCCTCCGCCA
redbury_ref-eIF4e 1 ATGGTTGTAGAAGACACCCCTAAATCCATCATCACCGACGATCAAATCACAGCAAACCCTAATAGGGTTATCGAAGACGACAACAATCTTGAAGAAGGAGAGATCCTCGATGAAGACGATTCCTCCGCCA

Lc_S_CN 105862 113 CTTCCAAACCCGTCGTCCACCAACCTCACCCCCTCGAGAATTCTTGGACTTTCTGGTTTGATACCCCTGCAGCAAAATCCAAACAAGCCGCTTGGGGTAGCTCAATGCGACCCATCTACACTTTTTCCAC
Lc_S_ILL 213AGL 131 CTTCCAAACCCGTCGTCCACCAACCTCACCCCCTCGAGAATTCTTGGACTTTCTGGTTTGATACCCCTGCAGCAAAATCCAAACAAGCCGCTTGGGGTAGCTCAATGCGACCCATCTACACTTTTTCCAC
Lc_S_ILL 2194 114 CTTCCAAACCCGTCGTCCACCAACCTCACCCCCTCGAGAATTCTTGGACTTTCTGGTTTGATACCCCTGCAGCAAAATCCAAACAAGCCGCTTGGGGTAGCTCAATGCGACCCATCTACACTTTTTCCAC
Lc_S_PI 329167 131 CTTCCAAACCCGTCGTCCACCAACCTCACCCCCTCGAGAATTCTTGGACTTTCTGGTTCGATACCCCTGCAGCAAAATCCAAACAAGCCGCTTGGGGTAGCTCAATGCGACCCATCTACACTTTTTCCAC
Lc_S_PI 472213 131 CTTCCAAACCCGTCGTCCACCAACCTCACCCCCTCGAGAATTCTTGGACTTTCTGGTTTGATACCCCTGCAGCAAAATCCAAACAAGCCGCTTGGGGTAGCTCAATGCGACCCATCTACACTTTTTCCAC
Lc_R_CDC Crimson 112 CTTCCAAACCCGTCGTCCACCAACCTCACCCCCTCGAGAATTCTTGGACTTTCTGGTTTGATACCCCTGCAGCAAAATCCAAACAAGCCGCTTGGGGTAGCTCAATGCGACCCATCTACACTTTTTCCAC
Lc_R_CDC Maxim 111 CTTCCAAACCCGTCGTCCACCAACCTCACCCCCTCGAGAATTCTTGGACTTTCTGGTTCGATACCCCTGCAGCAAAATCCGAACAAGCCGATTGGGGTAGCTCAATGCGACCCATCTACACTTTTTCCAC
Lc_R_CDC Greenstar 131 CTTCCAAACCCGTCGTCCACCAACCTCACCCCCTCGAGAATTCTTGGACTTTCTGGTTCGATACCCCTGCAGCAAAATCCAAACAAGCCGATTGGGGTAGCTCAATGCGACCCATCTACACTTTTTCCAC
Lc_R_CDC Impower 109 CTTCCAAACCCGTCGTCCACCAACCTCACCCCCTCGAGAATTCTTGGACTTTCTGGTTCGATACCCCTGCAGCAAAATCCAAACAAGCCGATTGGGGTAGCTCAATGCGACCCATCTACACTTTTTCCAC
Lc_R_PI 518734 131 CTTCCAAACCCGTCGTCCACCAACCTCACCCCCTCGAGAATTCTTGGACTTTCTGGTTCGATACCCCTGCAGCAAAATCCAAACAAGCCGATTGGGGTAGCTCAATGCGACCCATCTACACTTTTTCCAC
Lc_R_PI 297745 131 CTTCCAAACCCGTCGTCCACCAACCTCACCCCCTCGAGAATTCTTGGACTTTCTGGTTCGATACCCCTGCAGCAAAATCCAAACAAGCCGATTGGGGTAGCTCAATGCGACCCATCTACACTTTTTCCAC
Lc_R_PI 368648 114 CTTCCAAACCCGTCGTCCACCAACCTCACCCCCTCGAGAATTCTTGGACTTTCTGGTTCGATACCCCTGCAGCAAAATCCGAACAAGCCGATTGGGGTAGCTCAATGCGACCCATCTACACTTTTTCCAC
redbury_ref-eIF4e 131 CTTCCAAACCCGTCGTCCACCAACCTCACCCCCTCGAGAATTCTTGGACTTTCTGGTTCGATACCCCTGCAGCAAAATCCGAACAAGCCGATTGGGGTAGCTCAATGCGACCCATCTACACTTTTTCCAC

Lc_S_CN 105862 243 TGTTGAAGAGTTCTGGAGCATTTACAATAACATTCATCATCCTAGTAAGTTGGCTGTGGGAGCAGATTTCTATTGTTTCAAGCATAAGATTGAACCGAAATGGGAGGATCCCATCTGTGCTAATGGTGGG
Lc_S_ILL 213AGL 261 TGTTGAAGAGTTCTGGAGCATTTACAATAACATTCATCATCCTAGTAAGTTGGCTGTGGGAGCAGATTTCTATTGTTTCAAGCATAAGATTGAACCGAAATGGGAGGATCCCATCTGTGCTAATGGTGGG
Lc_S_ILL 2194 244 TGTTGAAGAGTTCTGGAGCATTTACAATAACATTCATCATCCTAGTAAGTTGGCTGTGGGAGCAGATTTCTATTGTTTCAAGCATAAGATTGAACCGAAATGGGAGGATCCCATCTGTGCTAATGGTGGG
Lc_S_PI 329167 261 TGTTGAAGAGTTCTGGAGCATTTACAATAACATTCATCATCCTAGTAAGTTGGCTGTGGGAGCAGATTTCTATTGTTTCAAGCATAAGATTGAACCGAAATGGGAGGATCCCATCTGTGCTAATGGTGGG
Lc_S_PI 472213 261 TGTTGAAGAGTTCTGGAGCATTTACAATAACATTCATCATCCTAGTAAGTTGGCTGTGGGAGCAGATTTCTATTGTTTCAAGCATAAGATTGAACCGAAATGGGAGGATCCCATCTGTGCTAATGGTGGG
Lc_R_CDC Crimson 242 TGTTGAAGAGTTCTGGAGCATTTACAATAACATTCATCATCCTAGTAAGTTGGCTGTGGGAGCAGATTTCTATTGTTTCAAGCATAAGATTGAACCGAAATGGGAGGATCCCATCTGTGCTAATGGTGGG
Lc_R_CDC Maxim 241 TGTTGAAGAGTTCTGGAGCATTTACAATAACATTCATCATCCTAGTAAGTTGGCTGTGAGAACAGATTTCTATTGCTTCAAGCATAAGATTAAACCGAAATGGGAGGATCCCATCTGTGCTAATGGTGGG
Lc_R_CDC Greenstar 261 TGTTGAAGAGTTCTGGAGCATTTACAATAACATTCATCATCCTAGTAAGTTGGCTGTGAGAACAGATTTCTATTGCTTCAAGCATAAGATTAAACCGAAATGGGAGGATCCCATCTGTGCTAATGGTGGG
Lc_R_CDC Impower 239 TGTTGAAGAGTTCTGGAGCATTTACAATAACATTCATCATCCTAGTAAGTTGGCTGTGAGAACAGATTTCTATTGCTTCAAGCATAAGATTAAACCGAAATGGGAGGATCCCATCTGTGCTAATGGTGGG
Lc_R_PI 518734 261 TGTTGAAGAGTTCTGGAGCATTTACAATAACATTCATCATCCTAGTAAGTTGGCTGTGAGAACAGATTTCTATTGCTTCAAGCATAAGATTAAACCGAAATGGGAGGATCCCATCTGTGCTAATGGTGGG
Lc_R_PI 297745 261 TGTTGAAGAGTTCTGGAGCATTTACAATAACATTCATCATCCTAGTAAGTTGGCTGTGAGAACAGATTTCTATTGCTTCAAGCATAAGATTAAACCGAAATGGGAGGATCCCATCTATGCTAATGGTGGG
Lc_R_PI 368648 244 TGTTGAAGAGTTCTGGAGCATTTACAATAACATTCATCATCCTAGTAAGTTGGCTGTGAGAACAGATTTCTATTGCTTCAAGCATAAGATTAAACCGAAATGGGAGGATCCCATCTATGCTAATGGTGGG
redbury_ref-eIF4e 261 TGTTGAAGAGTTCTGGAGCATTTACAATAACATTCATCATCCTAGTAAGTTGGCTGTGAGAACAGATTTCTATTGCTTCAAGCATAAGATTAAACCGAAATGGGAGGATCCCATCTGTGCTAATGGTGGG

Lc_S_CN 105862 373 AAATGGACTGCGAACTATCAGAAGGGAAAATCTGATACCAGTTGGTTGTATACGTTGTTGGCAATGATTGGAGAGCAATTTGATCATGGAGATGAAATTTGCGGAGCGGTTGTGAATGTAAGGGGTAGAG
Lc_S_ILL 213AGL 391 AAATGGACTGCGAACTATCAGAAGGGAAAATCTGATACCAGTTGGTTGTATACGTTGTTGGCAATGATTGGAGAGCAATTTGATCATGGAGATGAAATTTGCGGAGCGGTTGTGAATGTAAGGGGTAGAG
Lc_S_ILL 2194 374 AAATGGACTGCGAACTATCAGAAGGGAAAATCTGATACCAGTTGGTTGTATACGTTGTTGGCAATGATTGGAGAGCAATTTGATCATGGAGATGAAATTTGCGGAGCGGTTGTGAATGTAAGGGGTAGAG
Lc_S_PI 329167 391 AAATGGACTGCGAACTATCAGAAGGGAAAATCTGATACCAGTTGGTTGTATACGTTGTTGGCAATGATTGGAGAGCAATTTGATCATGGAGATGAAATTTGCGGAGCGGTTGTGAATGTAAGGGGTAGAG
Lc_S_PI 472213 391 AAATGGACTGCGAACTATCAGAAGGGAAAATCTGATACCAGTTGGTTGTATACGTTGTTGGCAATGATTGGAGAGCAATTTGATCATGGAGATGAAATTTGCGGAGCGGTTGTGAATGTAAGGGGTAGAG
Lc_R_CDC Crimson 372 AAATGGACTGCGAACTATCAGAAGGGAAAATCTGATACCAGTTGGTTGTATACGTTGTTGGCAATGATTGGAGAGCAATTTGATCATGGAGATGAAATTTGCGGAGCGGTTGTGAATGTAAGGGGTAGAG
Lc_R_CDC Maxim 371 AAATGGACTGCGAACTATCAGAAGGGAAAATCTGATACCAGTTGGTTGTATACGTTGTTGGCAATGATTGGAGAGCAATTTGATCATGGAGATGAAATTTGCGGAGCGGTTGTGAATGTAAGGGGTAGAG
Lc_R_CDC Greenstar 391 AAATGGACTGCGAACTATCAGAAGGGAAAATCTGATACCAGTTGGTTGTATACGTTGTTGGCAATGATTGGAGAGCAATTTGATCATGGAGATGAAATTTGCGGAGCGGTTGTGAATGTAAGGGGTAGAG
Lc_R_CDC Impower 369 AAATGGACTGCGAACTATCAGAAGGGAAAATCTGATACCAGTTGGTTGTATACGTTGTTGGCAATGATTGGAGAGCAATTTGATCATGGAGATGAAATTTGCGGAGCGGTTGTGAATGTAAGGGGTAGAG
Lc_R_PI 518734 391 AAATGGACTGCGAACTATCAGAAGGGAAAATCTGATACCAGTTGGTTGTATACGTTGTTGGCAATGATTGGAGAGCAATTTGATCATGGAGATGAAATTTGCGGAGCGGTTGTGAATGTAAGGGGTAGAG
Lc_R_PI 297745 391 AAATGGACTGCGAACTATCAGAAGGGAAAATCTGATACCAGTTGGTTGTATACGTTGTTGGCAATGATTGGAGAGCAATTTGATCATGGAGATGAAATTTGCGGAGCGGTTGTGAATGTAAGGGGTAGAG
Lc_R_PI 368648 374 AAATGGACTGCGAACTATCAGAAGGGAAAATCTGATACCAGTTGGTTGTATACGTTGTTGGCAATGATTGGAGAGCAATTTGATCATGGAGATGAAATTTGCGGAGCGGTTGTGAATGTAAGGGGTAGAG
redbury_ref-eIF4e 391 AAATGGACTGCGAACTATCAGAAGGGAAAATCTGATACCAGTTGGTTGTATACGTTGTTGGCAATGATTGGAGAGCAATTTGATCATGGAGATGAAATTTGCGGAGCGGTTGTGAATGTAAGGGGTAGAG

Lc_S_CN 105862 503 CTGAGAAAATTTCTATTTGGACTAAGAATGCTTCAAATGAAGCTGCTCAGGTGAGCATTGGAAAACAGTGGAAGGAGTTTCTTGATTATAATGAGACCATGGGCTTTATATTTCATGATGATGCAAGGAA
Lc_S_ILL 213AGL 521 CTGAGAAAATTTCTATTTGGACTAAGAATGCTTCAAATGAAGCTGCTCAGGTGAGCATTGGAAAACAGTGGAAGGAGTTTCTTGATTATAATGAGACCATGGGCTTTATATTTCATGATGATGCAAGGAA
Lc_S_ILL 2194 504 CTGAGAAAATTTCTATTTGGACTAAGAATGCTTCAAATGAAGCTGCTCAGGTGAGCATTGGAAAACAGTGGAAGGAGTTTCTTGATTATAATGAGACCATGGGCTTTATATTTCATGATGATGCAAGGAA
Lc_S_PI 329167 521 CTGAGAAAATTTCTATTTGGACTAAGAATGCTTCAAATGAAGCTGCTCAGGTGAGCATTGGAAAACAGTGGAAGGAGTTTCTTGATTATAATGAGACCATGGGCTTTATATTTCATGATGATGCAAGGAA
Lc_S_PI 472213 521 CTGAGAAAATTTCTATTTGGACTAAGAATGCTTCAAATGAAGCTGCTCAGGTGAGCATTGGAAAACAGTGGAAGGAGTTTCTTGATTATAATGAGACCATGGGCTTTATATTTCATGATGATGCAAGGAA
Lc_R_CDC Crimson 502 CTGAGAAAATTTCTATTTGGACTAAGAATGCTTCAAATGAAGCTGCTCAGGTGAGCATTGGAAAACAGTGGAAGGAGTTTCTTGATTATAATGAGACCATGGGCTTTATATTTCATGATGATGCAAGGAA
Lc_R_CDC Maxim 501 CTGAGAAAATTTCTATTTGGACTAAGAATGCTTCAAATGAAGCTGCTCAGGTGAGCATTGGAAAACAGTGGAAGGAGTTTCTTGATTATAATGAGACCATGGGCTTTATATTTCATGATGATGCAAGGAA
Lc_R_CDC Greenstar 521 CTGAGAAAATTTCTATTTGGACTAAGAATGCTTCAAATGAAGCTGCTCAGGTGAGCATTGGAAAACAGTGGAAGGAGTTTCTTGATTATAATGAGACCATGGGCTTTATATTTCATGATGATGCAAGGAA
Lc_R_CDC Impower 499 CTGAGAAAATTTCTATTTGGACTAAGAATGCTTCAAATGAAGCTGCTCAGGTGAGCATTGGAAAACAGTGGAAGGAGTTTCTTGATTATAATGAGACCATGGGCTTTATATTTCATGATGATGCAAGGAA
Lc_R_PI 518734 521 CTGAGAAAATTTCTATTTGGACTAAGAATGCTTCAAATGAAGCTGCTCAGGTGAGCATTGGAAAACAGTGGAAGGAGTTTCTTGATTATAATGAGACCATGGGCTTTATATTTCATGATGATGCAAGGAA
Lc_R_PI 297745 521 CTGAGAGAATTTCTATTTGGACTAAGAATGCTTCAAATGAAGCTGCTCAGGTGAGCATTGGAAAACAGTGGAAGGAGTTTCTTGATTATAATGAGACCATGGGCTTTATATTTCATGATGATGCAAGGAA
Lc_R_PI 368648 504 CTGAGAAAATTTCTATTTGGACTAAGAATGCTTCAAATGAAGCTGCTCAGGTGAGCATTGGAAAACAGTGGAAGGAGTTTCTTGATTATAATGAGACCATGGGCTTTATATTTCATGATGATGCAAGGAA
redbury_ref-eIF4e 521 CTGAGAAAATTTCTATTTGGACTAAGAATGCTTCAAATGAAGCTGCTCAGGTGAGCATTGGAAAACAGTGGAAGGAGTTTCTTGATTATAATGAGACCATGGGCTTTATATTTCATGATGATGCAAGGAA

Lentil eukaryotic translation initiation factor 4E (*LceIF4e*) nucleotide sequence alignment of 12 lentil lines from the study in reference to CDC redberry (Ramsay et al. 2021). The sequence of 12 lines was generated from sanger sequencing. Full eIF4E coding sequence can be found in NCBI with accession number as follow: CDC Greenstar (PZ761244), CDC Impower (PZ761245), CDC Maxim (PZ761246), Crimson AGL (PZ761247), PI 297745 (PZ761248), PI 368648 (PZ761249), PI 518734 (PZ761250), CN105862 (PZ761251), ILL 213_AGL (PZ761252), ILL 2194 (PZ761253), PI 329167 (PZ761254), PI 472213 (PZ761255). 

```
Lc_S_CN105862      1   ------PKSIITDDQITANPNRVIEDDNNLEEGEILDEDDSSATSKPVVHQPHPLENSWTFWFDTPAAKSKQAAWGSSMRPIYTFSTVEEFWSIYNNIHH
Lc_S_ILL213AGL     1   MVVEDTPKSIITDDQITANPNRVIEDDNNLEEGEILDEDDSSATSKPVVHQPHPLENSWTFWFDTPAAKSKQAAWGSSMRPIYTFSTVEEFWSIYNNIHH
Lc_S_PI329167      1   MVVEDTPKSIITDDQITANPNRVIEDDNNLEEGEILDEDDSSATSKPVVHQPHPLENSWTFWFDTPAAKSKQAAWGSSMRPIYTFSTVEEFWSIYNNIHH
Lc_S_ILL2194       1   ------PKSIITDDQITANPNRVIEDDNNLEEGEILDEDDSSATSKPVVHQPHPLENSWTFWFDTPAAKSKQAAWGSSMRPIYTFSTVEEFWSIYNNIHH
Lc_S_PI472213      1   MVVEDTPKSIITDDQITANPNRVIEDDNNLEEGEILDEDDSSATSKPVVHQPHPLENSWTFWFDTPAAKSKQAAWGSSMRPIYTFSTVEEFWSIYNNIHH
Lc_R_CrimsonAGL    1   -------KSIITDDQITANPNRVIEDDNNLEEGEILDEDDSSATSKPVVHQPHPLENSWTFWFDTPAAKSKQAAWGSSMRPIYTFSTVEEFWSIYNNIHH
Lc_R_CDC-Maxim     1   -------KSIITDDQITANPNRVIEDDNNLEEGEILDEDDSSATSKPVVHQPHPLENSWTFWFDTPAAKSEQADWGSSMRPIYTFSTVEEFWSIYNNIHH
Lc_R_CDC-Greenstar 1   MVVEDTPKSIITDDQITANPNRVIEDDNNLEEGEILDEDDSSATSKPVVHQPHPLENSWTFWFDTPAAKSKQADWGSSMRPIYTFSTVEEFWSIYNNIHH
Lc_R_CDC-Impower   1   --------SIITDDQITANPNRVIEDDNNLEEGEILDEDDSSATSKPVVHQPHPLENSWTFWFDTPAAKSKQADWGSSMRPIYTFSTVEEFWSIYNNIHH
Lc_R_PI518734      1   MVVEDTPKSIITDDQITANPNRVIEDDNNLEEGEILDEDDSSATSKPVVHQPHPLENSWTFWFDTPAAKSKQADWGSSMRPIYTFSTVEEFWSIYNNIHH
Lc_R_PI297745      1   MVVEDTPKSIITDDQITANPNRVIEDDNNLEEGEILDEDDSSATSKPVVHQPHPLENSWTFWFDTPAAKSKQADWGSSMRPIYTFSTVEEFWSIYNNIHH
Lc_R_PI368648      1   ------PKSIITDDQITANPNRVIEDDNNLEEGEILDEDDSSATSKPVVHQPHPLENSWTFWFDTPAAKSEQADWGSSMRPIYTFSTVEEFWSIYNNIHH
Lc_ref_redberry    1   MVVEDTPKSIITDDQITANPNRVIEDDNNLEEGEILDEDDSSATSKPVVHQPHPLENSWTFWFDTPAAKSEQADWGSSMRPIYTFSTVEEFWSIYNNIHH
Ps_R_PI347464      1   MVVEDTPKSIITDDQITTNPNRVIEDDNNLEEGEILDEDDSSATSKPVVHQPHLLENSWTFLFDTPAAKSKQDDWGSSMRPIYTFSTVEEFWSIYNNIHH
Ps_R_PI193835      1   MVVEDTPKSIITDDQITTNPNRVIEDDNNLEEGEILDEDDSSATSKPVVHQPHLLENSWTFLFDTPAAKSKQDDWGSSMRPIYTFSTVEEFWSIYNNIHH
Ps_R_PI269818      1   MVVEETPKSIITDDQITTNPNRVIEDDNNLEEGEILDEDDSSATSKPVVHQPHLLENSWTFWFDTPAAKSKQPDWGS-MRPIYTFSTVEEFWSIYNNIHH
Ps_R_JI1405        1   MVVEETPKSIITDDQITTNPNRVIEDDNNLEEGEILDEDDSSATSKPVVHQPHLLENSWTFLFDTPAAKSKQDDWGSSMRPIYTFSTVEEFWSIYNNIHH
Ps_S_DSP           1   MVVEDTPKSIITDDQITTNPNRVIEDDNNLEEGEILDEDDSSATSKPVVHQPHLLENSWTFWFDTPAAKSKQAAWGSSMRPIYTFSTVEEFWSIYNNIHH
Ps_S_Brutus        1   MVVEDTPKSIITDDQITTNPNRVIEDDNNLEEGEILDEDDSSATSKPVVHQPHLLESSWTFWFDTPAAKSKQAAWGSSMRPIYTFSTVEEFWSIYNNIHH
Ps_S_Bonneville    1   MVVEDTPKSIITDDQITTNPNRVIEDDNNLEEGEILDEDDSSATSKPVVHQPHLLENPWTFWFDTPAAKSKQAAWGSSMRPIYTFSTVEEFWSIYNNIHH
Ps_S_Fjord         1   MVVEDTPKSIITDDQITTNPNRVIEDDNNLEEGEILDEDDSSATSKPVVHQPHLLENSWTFWFDTPAAKSKQAAWGSSMRPIYTFSTVEEFWSIYNNIHH

Lc_S_CN105862      95  PSKLAVGADFYCFKHKIEPKWEDPICANGGKWTANYQKGKSDTSWLYTLLAMIGEQFDHGDEICGAVVNVRGRAEKISIWTKNASNEAAQVSIGKQWKEF
Lc_S_ILL213AGL     101 PSKLAVGADFYCFKHKIEPKWEDPICANGGKWTANYQKGKSDTSWLYTLLAMIGEQFDHGDEICGAVVNVRGRAEKISIWTKNASNEAAQVSIGKQWKEF
Lc_S_PI329167      101 PSKLAVGADFYCFKHKIEPKWEDPICANGGKWTANYQKGKSDTSWLYTLLAMIGEQFDHGDEICGAVVNVRGRAEKISIWTKNASNEAAQVSIGKQWKEF
Lc_S_ILL2194       95  PSKLAVGADFYCFKHKIEPKWEDPICANGGKWTANYQKGKSDTSWLYTLLAMIGEQFDHGDEICGAVVNVRGRAEKISIWTKNASNEAAQVSIGKQWKEF
Lc_S_PI472213      101 PSKLAVGADFYCFKHKIEPKWEDPICANGGKWTANYQKGKSDTSWLYTLLAMIGEQFDHGDEICGAVVNVRGRAEKISIWTKNASNEAAQVSIGKQWKEF
Lc_R_CrimsonAGL    94  PSKLAVGADFYCFKHKIEPKWEDPICANGGKWTANYQKGKSDTSWLYTLLAMIGEQFDHGDEICGAVVNVRGRAEKISIWTKNASNEAAQVSIGKQWKEF
Lc_R_CDC-Maxim     94  PSKLAVRTDFYCFKHKIKPKWEDPICANGGKWTANYQKGKSDTSWLYTLLAMIGEQFDHGDEICGAVVNVRGRAEKISIWTKNASNEAAQVSIGKQWKEF
Lc_R_CDC-Greenstar 101 PSKLAVRTDFYCFKHKIKPKWEDPICANGGKWTANYQKGKSDTSWLYTLLAMIGEQFDHGDEICGAVVNVRGRAEKISIWTKNASNEAAQVSIGKQWKEF
Lc_R_CDC-Impower   93  PSKLAVRTDFYCFKHKIKPKWEDPICANGGKWTANYQKGKSDTSWLYTLLAMIGEQFDHGDEICGAVVNVRGRAEKISIWTKNASNEAAQVSIGKQWKEF
Lc_R_PI518734      101 PSKLAVRTDFYCFKHKIKPKWEDPICANGGKWTANYQKGKSDTSWLYTLLAMIGEQFDHGDEICGAVVNVRGRAEKISIWTKNASNEAAQVSIGKQWKEF
Lc_R_PI297745      101 PSKLAVRTDFYCFKHKIKPKWEDPIYANGGKWTANYQKGKSDTSWLYTLLAMIGEQFDHGDEICGAVVNVRGRAERISIWTKNASNEAAQVSIGKQWKEF
Lc_R_PI368648      95  PSKLAVRTDFYCFKHKIKPKWEDPIYANGGKWTANYQKGKSDTSWLYTLLAMIGEQFDHGDEICGAVVNVRGRAEKISIWTKNASNEAAQVSIGKQWKEF
Lc_ref_redberry    101 PSKLAVRTDFYCFKHKIKPKWEDPICANGGKWTANYQKGKSDTSWLYTLLAMIGEQFDHGDEICGAVVNVRGRAEKISIWTKNASNEAAQVSIGKQWKEF
Ps_R_PI193835      101 PGKLAVRADFYCFKHKIEPKWEDPICANGGKWTANYPKGKSDTSWLYTLLAMIGEQFDHGDEICGAVVKVRGRAEKISIWTKNASNEAAQVSIGKQWKEF
Ps_R_PI193835      101 PGKLAVRADFYCFKHKIEPKWEDPICANGGKWTANYPKGKSDTSWLYTLLAMIGEQFDHGDEICGAVVKVRGRAEKISIWTKNASNEAAQVSIGKQWKEF
Ps_R_PI269818      100 PGKLAVGADFYCFKHKIEPKWEDPICANGGKWTANYPKGKSDTSWLYTLLAMIGEQFDHGDEICGAVVNVRGRAEKISIWTKNASNEAAQVSIGKQWKEF
Ps_R_JI1405        101 PGKLAVRADFYCFKHKIEPKWEDPICANGGKWTANYPKGKSDTSWLYTLLAMIGEQFDHGDEICGAVVKVRGRAEKISIWTKNASNEAAQVSIGKQWKEF
Ps_S_DSP           101 PGKLAAGADFYCFKHKIEPKWEDPICANGGKWTANYPKGKSDTSWLYTSLAMIGEQFDHGDEICGAVVNVRGRAEKISIWTKNASNEAAQVSIGKQWKEF
Ps_S_Brutus        101 PGKLAVGADFYCFKHKIEPKWEDPICANGGKWTANYPKGKSDTSWLYTLLAMIGEQFDHGDEICGAVVNVGGRAEKISIWTKNASNEAAQVSIGKQWKEF
Ps_S_Bonneville    101 PGKLAVGADFYCFKHKIEPKWEDPICANGGKWTANYPKGKSDTSWLYTLLAMIGEQFDHGDEICGAVVNVRGRAEKISIWTKNASNEAAQVSIGKQWKEF
Ps_S_Fjord         101 PGKLAVGADFYCFKHKIEPKWEDPICANGGKWTANYPKGKSDTSWLYTLLAMIGEQFDHGDEICGAVVNVRGRAEKISIWTKNASNEAAQVSIGKQWKEF


```

Amino acid sequence alignment of eukaryotic translation initiation factor 4E (eIF4E) from 13 lentil (Lc) and 8 pea (Ps) lines. PSbMV susceptible lines are denoted with S (e.g. Ps_**S**_Fjord) and the resistance lines are denoted with R (e.g. Ps_**S**_DSP). The amino acid sequences of all lentils were generated by translating the nucleotide sequence (supplementary file 1) in SnapGene (v8.2.2). The NCBI accession numbers of the aligned pea amino acid sequences are as follows: CDC Greenstar (YGE40059.1), CDC Impower (YGE40060.1), CDC Maxim (YGE40061.1), Crimson AGL (YGE40062.1), PI 297745 (YGE40063.1), PI 368648 (YGE40064.1), PI 518734 (YGE40065.1), CN105862 (YGE40066.1), ILL 213_AGL (YGE40067.1), ILL 2194 (YGE40068.1), PI 329167 (YGE40069.1), PI 472213 (YGE40070.1). Brutus (DQ641470), Bonneville (DQ641471), Fjord (GU289734), DSP (DQ641472), PI347464 (DQ641473), PI193835 (DQ641474), PI269818 (AY611425), JI1405 (AY611423).

```
CDC KR-1 AGL_1      1   AATCCATCATCACCGACGATCAAATCACAGCAAACCCTAATAGGGTTATCGAAGACGACAACAATCTTGAAGAAGGAGAGATCCTCGATGAAG
CDC KR-1 AGL_2      1   AATCCATCATCACCGACGATCAAATCACAGCAAACCCTAATCGGGTTATCGAAGACGACAACAATCTTGAAGAAGGAGAGATCCTCGATGAAG
ILL 4665 AGL_1      1   AATCCATCATCACCGACGATCAAATCACAGCAAACCCTAATAGGGTTATCGAAGACGACAACAATCTTGAAGAAGGAGAGATCCTCGATGAAG
ILL 4665 AGL_2      1   AATCCATCATCACCGACGATCAAATCACAGCAAACCCTAATCGGGTTATCGAAGACGACAACAATCTTGAAGAAGGAGAGATCCTCGATGAAG
PI 431893 LSP AGL_1 1   AATCCATCATCACCGACGATCAAATCACAGCAAACCCTAATAGGGTTATCGAAGACGACAACAATCTTGAAGAAGGAGAGATCCTCGATGAAG
PI 431893 LSP AGL_2 1   AATCCATCATCACCGACGATCAAATCACAGCAAACCCTAATCGGGTTATCGAAGACGACAACAATCTTGAAGAAGGAGAGATCCTCGATGAAG

CDC KR-1 AGL_1      94  ACGATTCCTCCGCCACTTCCAAACCCGTCGTCCACCAACCTCACCCCCTCGAGAATTCTTGGACTTTCTGGTTCGATACCCCTGCAGCAAAATCCGAACAAGCCGATTGG
CDC KR-1 AGL_2      94  ACGATTCCTCCGCCACTTCCAAACCCGTCGTCCACCAACCTCACCCCCTCGAGAATTCTTGGACTTTCTGGTTTGATACCCCTGCAGCAAAATCCAAACAAGCCGCTTGG
ILL 4665 AGL_1      94  ACGATTCCTCCGCCACTTCCAAACCCGTCGTCCACCAACCTCACCCCCTCGAGAATTCTTGGACTTTCTGGTTCGATACCCCTGCAGCAAAATCCAAACAAGCCGATTGG
ILL 4665 AGL_2      94  ACGATTCCTCCGCCACTTCCAAACCCGTCGTCCACCAACCTCACCCCCTCGAGAATTCTTGGACTTTCTGGTTTGATACCCCCGCAGCAAAATCCAAACAAGCCGCTTGG
PI 431893 LSP AGL_1 94  ACGATTCCTCCGCCACTTCCAAACCCGTCGTCCACCAACCTCACCCCCTCGAGAATTCTTGGACTTTCTGGTTCGATACCCCTGCAGCAAAATCCAAACAAGCCGATTGG
PI 431893 LSP AGL_2 94  ACGATTCCTCCGCCACTTCCAAACCCGTCGTCCACCAACCTCACCCCCTCGAGAATTCTCGGACTTTCTGGTTCGATACCCCCGCAGCAAAATCCAAACAAGCCGATTGG

CDC KR-1 AGL_1      204 GGTAGCTCAATGCGACCCATCTACACTTTTTCCACTGTTGAAGAGTTCTGGAGCATTTACAATAACATTCATCATCCTAGTAAGTTGGCTGTGAGAACAGATTTCTATTG
CDC KR-1 AGL_2      204 GGTAGCTCAATGCGACCCATCTACACTTTTTCCACTGTTGAAGAGTTCTGGAGCATTTACAATAACATTCATCATCCTAGTAAGTTGGCTGTGGGAGCAGATTTCTATTG
ILL 4665 AGL_1      204 GGTAGCTCAATGCGACCCATCTACACTTTTTCCACTGTTGAAGAGTTCTGGAGCATTTACAATAACATTCATCATCCTAGTAAGTTGGCTGTGAGAACAGATTTCTATTG
ILL 4665 AGL_2      204 GGTAGCTCAATGCGACCCATCTACACTTTTTCCACTGTTGAAGAGTTCTGGAGCATTTACAATAACATTCATCATCCTAGTAAGTTGGCTGTGGGAGCAGATTTCTATTG
PI 431893 LSP AGL_1 204 GGTAGCTCAATGCGACCCATCTACACTTTTTCCACTGTTGAAGAGTTCTGGAGCATTTACAATAACATTCATCATCCTAGTAAGTTGGCTGTGAGAACAGATTTCTATTG
PI 431893 LSP AGL_2 204 GGTAGCTCAATGCGACCCATCTACACTTTTTCCACTGTTGAAGAGTTCTGGAGCATTTACAATAACATTCATCATCCTAGTAAGTTGGCTGTGGGAGCAGATTTCTATTG

CDC KR-1 AGL_1      314 CTTCAAGCATAAGATTAAACCGAAATGGGAGGATCCCATCTGTGCTAATGGTGGG
CDC KR-1 AGL_2      314 TTTCAAGCATAAGATTGAACCGAAATGGGAGGATCCCATCTGTGCTAATGGTGGG
ILL 4665 AGL_1      314 CTTCAAGCATAAGATTAAACCGAAATGGGAGGATCCCATCTGTGCTAATGGTGGG
ILL 4665 AGL_2      313 TTTCAAGCATAAGATTGAACCGAAATGGGAGGATCCCATCTGTGCTAATGGTGGG
PI 431893 LSP AGL_1 314 CTTCAAGCATAAGATTAAACCGAAATGGGAGGATCCCATCTGTGCTAATGGTGGG
PI 431893 LSP AGL_2 314 TTTCAAGCATAAGATTGAACCGAAATGGGAGGATCCCATCTGTGCTAATGGTGGG

```

Lentil eukaryotic translation initiation factor 4E (*LceIF4e*) nucleotide sequence alignment of 6 samples from three lentil lines showing heterogeneous allele call in one of the samples.
